# The phosphatases TCPTP, PTPN22, and SHP1 play unique roles in T cell phosphotyrosine maintenance and feedback regulation of the TCR

**DOI:** 10.1101/2025.05.05.652301

**Authors:** Aurora Callahan, Aisharja Mojumdar, Mengzhou Hu, Amber Wang, Alijah A Griffith, Nicholas Huang, Xien Yu Chua, Nick Mroz, Ryan Z. Puterbaugh, Shanelle P Reilly, Arthur R. Salomon

## Abstract

The protein tyrosine phosphatases (PTPs) TCPTP, PTPN22, and SHP1 are critical regulators of the activating phosphotyrosine (pY) site on the initiating T cell kinase, Lck^Y394. Still,^ the broader implications of these phosphatases in T cell receptor (TCR) signalling and T cell biology remain unclear. By combining CRISPR/Cas9 gene editing and mass spectrometry, we evaluate the protein- and pY-level effects of TCPTP, PTPN22, and SHP1 in the Jurkat T cell model system. We find that deletion of each phosphatase corresponds to unique changes in the proteome of T cells, with few large-scale changes to TCR signalling proteins. Notably, PTPN22 and SHP1 deletions have opposing effects on pY abundance globally, while TCPTP deletion modestly elevates pY levels. Finally, we show that TCPTP is indirectly involved in Erk1/2 positive feedback to the TCR. Overall, our work provides evidence for alternative functions of three T cell phosphatases long thought to be redundant.

## Introduction

T cell activation is a delicate signalling process, and mutations that affect the timing or strength of T cell activation often result in autoimmune disorders^1,2^. T cell activation is achieved through binding of a peptide-loaded major histocompatibility complex (pMHC) to the α - and β-subunits of the T cell receptor (TCR), promoting dephosphorylation of the Src-family tyrosine kinase Lck at tyrosine 505 (Lck^Y505^) and subsequent autophosphorylation and activation of Lck^Y394^. Lck then phosphorylates immunoreceptor tyrosine-based activation motifs (ITAMs) on TCR subunits CD3ε/δ/γ, which act as a docking site for the Syk-family kinase Zap70. After Zap70 phosphorylation and activation by Lck, Lck and Zap70 phosphorylate the scaffolding proteins LAT and SLP76 and their associated binding partners, forming the TCR signalosome. Signal diversifies in the TCR signalosome to coordinate gene expression, rearrangement of the actin cytoskeleton, and release of proteins with immune system activating or cytotoxic function (T cell signalling has been reviewed extensively in Gaud et al.2018^1^). Despite a large body of literature describing the feed-forward activation of T cells from TCR/pMHC binding, much less is known about the feedback mechanisms involved in maintaining homeostasis and regulation along the TCR signalling pathway.

Our current understanding of TCR signalling regulation focuses on the initiating kinase, Lck, and three phosphorylation sites known to play a role in activation or substrate specificity: Lck^Y505^, Lck^Y394^, and Lck^Y192^. When phosphorylated by Csk, Lck^Y505^ locks Lck in an inactive conformation, preventing Lck^Y394^ autophosphorylation and Lck activation. Thus, its dephosphorylation by CD45 is a necessary step in Lck activation^3-5^. Phosphorylation of Lck^Y192^ is known to prevent Lck/CD45 interactions, promote interactions with Zap70, TCRζ, and PLCγ1, and regulate the PLCγ1-dependent activation of Erk1/2^6–9^. Phosphorylation of Lck^Y394^ increases its kinase activity, which can be dampened by dephosphorylation. Five protein tyrosine phosphatases (PTPs) are known to dephosphorylate Lck^Y394^: CD45, T cell protein tyrosine phosphatase (TCPTP), PTP non-receptor type 22 (PTPN22), SH2 domain-containing tyrosine phosphatase 1 (SHP1), and JNK-associated phosphatase (JKAP)^10-16^. The recruitment of TCPTP and PTPN22 to Lck and the early T cell receptor components is ambiguous, with no known recruitment strategy for TCPTP and conflicting literature for CSK-dependent and TRAF3-dependent recruitment of PTPN22^17–19^. Thus, the biological significance of TCPTP and PTPN22 dephosphorylation of Lck^Y394^ remains unclear. In contrast, JKAP is anchored in the plasma membrane by myristoylation, similarly to Lck, which is thought to be necessary for JKAP-dependent dephosphorylation of Lck^Y394 14,20^.

Lastly, SHP1 can bind directly to the C-terminal region of Lck, which is disrupted by Lck^S59^ phosphorylation by Erk1/2. However, SHP1 also associates with GRB2 and GRB2-associated proteins via SH2 domain-mediated binding to shared or adjacent phosphotyrosine-containing motifs on adapter proteins such as LAT^21–23^. Regardless, TCPTP, PTPN22, SHP1, and JKAP dysregulation are implicated in numerous autoimmune disorders for their role in maintaining T cells’ homeostasis^2,14,24–44^.

Our laboratory and others have previously defined a positive feedback mechanism connecting Erk1/2 to TCR signal transduction^21^. Loss of Erk1/2 activation through small molecule inhibition of MEK1/2 leads to a global decrease in tyrosine phosphorylation, which mutes TCR signalling responses to soluble antibody or pMHC ligation^21,23,45^. The leading hypothesis, in which Erk1/2 phosphorylates Lck^S59^ to prevent SHP1 docking and Lck^Y394^ dephosphorylation, remains ambiguous as Lck^S59A^ knock-in mice fail to show overt defects in T cell activation and ligand discrimination^23^. Due to the redundancy of Lck^Y394^ dephosphorylation, we hypothesise that Lck-regulating PTPs may be involved in Erk1/2 positive feedback. Using the Jurkat T cell model system, CRISPR/Cas9 gene editing, and mass spectrometry-based proteomics, we evaluate the roles of TCPTP, PTPN22, and SHP1 in maintaining proteomic homeostasis, maintaining pY homeostasis, and Erk1/2 positive feedback. We find that T cell signalling proteins were largely unaffected by the knockout of each phosphatase. However, J.PTPN22 shows significant dehancement of KEGG T cell signalling proteins. TCPTP, PTPN22, and SHP1 play opposing roles in pY maintenance globally, with PTPN22 negatively regulating, SHP1 positively regulating, and TCPTP modestly negatively regulating pY abundance. Finally, while inhibition of MEK1/2 with U0126 reduces tyrosine phosphorylation globally irrespective of PTP KO, TCPTP KO is affected minimally by U0126 and recovered quickly after soluble antibody stimulation of the TCR, suggesting a role for TCPTP in Erk positive feedback.

## Results

### *PTPN2* (TCPTP), *PTPN22* (PTPN22), and *PTPN6* (SHP1) knockouts do not affect the abundance or retention of key TCR signalling proteins

To evaluate the specific effects of TCPTP, PTPN22, and SHP1 on T cell homeostasis and feedback regulation of the TCR, we generated genomic KOs of *PTPN2*, *PTPN22*, and *PTPN6* in Jurkat T cells using CRISPR/Cas9-based gene editing (Figure 1A). Clonal KOs were screened for TCPTP, PTPN22, and SHP1 expression by Western blot, and one clone was selected for use (Figure 1A-B). The resulting clones, J.TCPTP-(T2 1C10), J.PTPN22-(P1 1C1), and J.SHP1-(S1 2C4) displayed surface levels of CD3ε and CD45 similar to WT Jurkats (clone E6.1), while J.SHP1- and J.TCPTP-showed varying levels of surface TCR□ by flow cytometry (Figure 1C). Using LC-MS/MS-based protein profiling, we evaluated the influence of *PTPN2*, *PTPN22*, and *PTPN6* knockout on global protein expression. We sequenced the proteome of each cell line deeply and reproducibly, with few imputed values per replicate, cell line-dependent clustering, and high linearity (Figure 1D, Supporting Figure 1A). Two samples, J.TCPTP-R5 and J.PTPN22-R4, show markedly reduced protein abundance compared with the other replicates (Supporting Figure 1B) and were removed from further analysis. We found that genomic deletions of *PTPN2* and *PTPN22* had a minimal effect on global protein abundance (Figures 1D-E), with few T cell signalling proteins showing significantly reduced abundance compared to WT Jurkats (Figure 1F).

**Figure 1:**
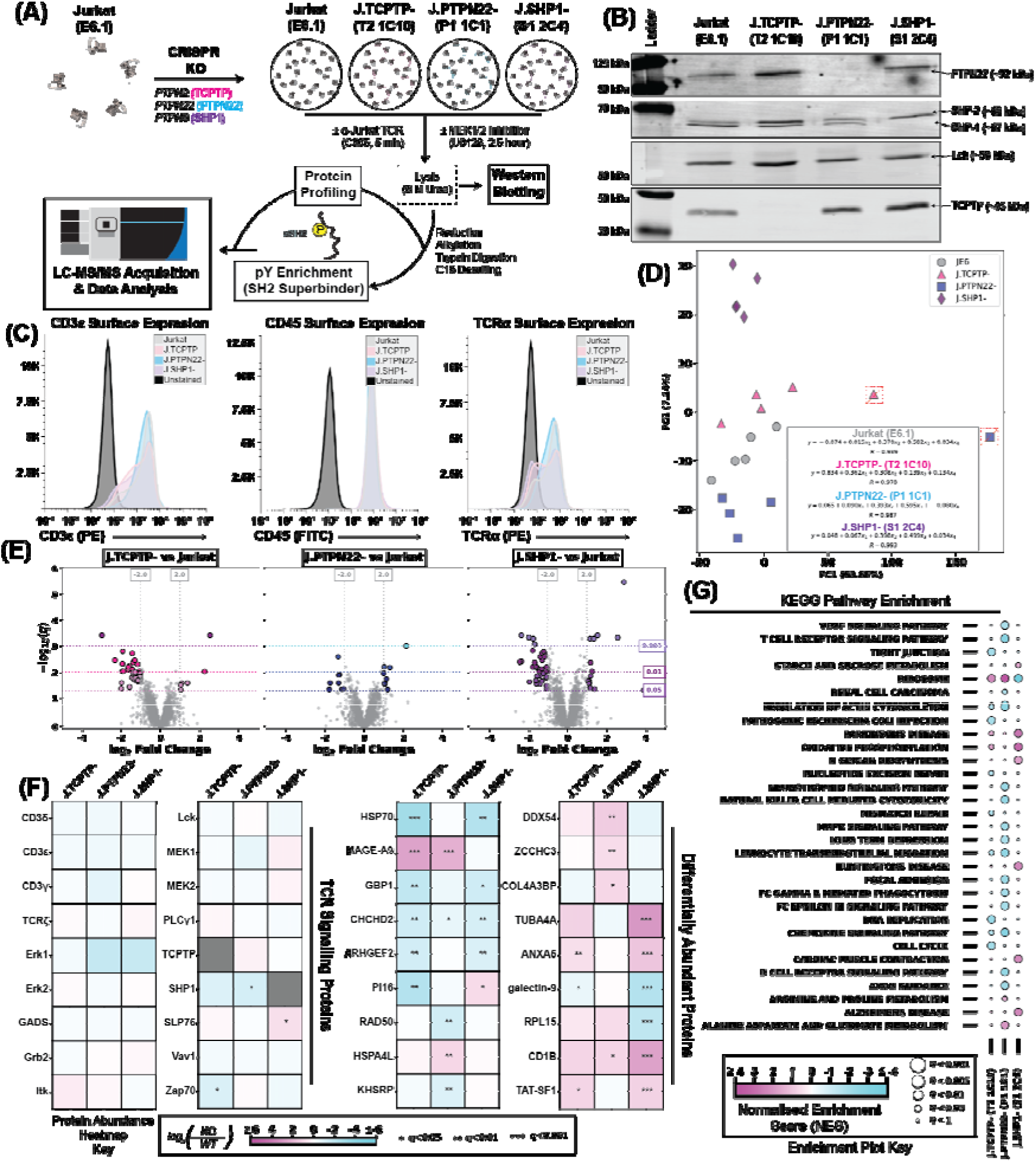
Generation and proteomic characterisation of J.TCPTP-, J.PTPN22-, and J.SHP1-. (A) Schematic representation of CRISPR Cas9 gene editing of Jurkat T cells and subsequent signalling analysis. (B) Western blot showing loss of PTPN22 (row 1), SHP1 (row 2), and TCPTP (row 4) in the selected clones. (C) Flow cytometry analysis of CD3ε (left), CD45 (middle), and TCRα (right) surface expression in the phosphatase KO clones. (D) Principal component analysis of LC-MS/MS protein profiling data for the selected clones. J.TCPTP-Replicate 5 and J.PTPN22- Replicate 4 are discarded due to sample quality (See Supporting Figure 1). (E) Volcano plot analysis showing differentially abundant proteins in J.TCPTP- (left), J.PTPN22- (middle), and J.SHP1- (right) relative to Jurkats. (F) Heatmaps showing specific protein abundance for T cell proteins (left) and differentially abundant proteins (right). (G) Single sample gene set enrichment analysis using the KEGG database for differentially abundant proteins.

Interestingly, *PTPN2* and *PTPN22* deletion increased the abundance of MAGE-A9, a protein of completely unknown function implicated in several cancers^46–49^. In contrast, *PTPN6* deletion broadly influenced non-TCR protein expression, including upregulation of TUBA4A and CD1b, and downregulation of galectin-9 and RPL15 (Figures 1E-F). While single-sample gene set enrichment analysis (ssGSEA) using the Kyoto Encyclopedia of Genes and Genomes (KEGG) pathway database^50^ revealed a significant dehancement of TCR signalling proteins in J.PTPN22-(Figure 1G), individual TCR signalling proteins were not significantly altered. In contrast, J.TCPTP-showed dehancement of Cell Cycle and DNA Replication pathways, and J.SHP1-showed enhancement of multiple metabolic pathways and disease-associated pathways (Figure 1G). Together, our data show that *PTPN2*, *PTPN22*, and *PTPN6* knockouts are successful and do not broadly impact individual proteins in the TCR pathway. However, they do appear to have specific effects on the proteome.

### *PTPN2* (TCPTP), *PTPN22* (PTPN22), and *PTPN6* (SHP1) knockouts respond to TCR stimulation and, to some degree, Erk positive feedback

To determine whether the knockouts remained responsive to T cell receptor stimulation and Erk positive feedback, we inhibited MEK1/2 using U0126 and stimulated cells with α-Jurkat TCR (clone C305) followed by Western blot analysis of various pY sites known to be TCR responsive (Figure 2A). Titration of U0126 from 0.2 μM to 50 μM showed competent Erk1^T202Y204^/Erk2^T185Y187^ phosphorylation until a U0126 concentration of 20 μM (Figures 2B-C, Supporting Figure 2), as seen previously^21^. Positive feedback from Erk1/2 to the TCR is well described and includes a phenotype of global pY depletion^21^. Using α-pY (clone 4G10), an antibody that robustly detects tyrosine phosphorylated T cell proteins^51^, we visualised changes in a small subset of pY proteins during TCR stimulation and U0126 treatment. We observed that U0126 reduced pY levels to the untreated basal state at 20 μM in all cell lines (Figure 2D, Supporting Figure 3). Since 20 μM U0126 appeared to completely ablate Erk1^T202Y204^/Erk2^T185Y187^ phosphorylation, we evaluated any changes in the kinetics of TCR signalling proteins with or without 20 μM U0126 treatment. As expected, Erk1^T202Y204^/Erk2^T185Y187^ phosphorylation was significantly reduced with U0126 treatment in all cell lines at all time points (Figures 2E-F, Supporting Figure 4). J.TCPTP-showed the highest recovery of Erk1^T202Y204^/Erk2^T185Y187^ phosphorylation during U0126 treatment of all cell lines after 5 minutes of TCR stimulation (p=0.0016 vs 0 min), with all lines except J.SHP1-showing significant Erk1^T202Y204^/Erk2^T185Y187^ phosphorylation after 10 minutes (Figure 2E-F, Supporting Figure 4). All cell lines except J.PTPN22-showed significant pY induction after 2 and 5 minutes of TCR stimulation (p < 0.05), and no cell line showed significant global pY induction after U0126 treatment despite clear increases in specific pY bands in J.TCPTP- and J.SHP1- (Figure 2G, Supporting Figures 5-6). Similarly, phosphorylation of the Src family kinase activation site (peptide NEpYTAR) at a molecular weight consistent with Lck (Supporting Figure 7) was induced at 2 minutes (p<0.05; Jurkat, J.TCPTP-, J.SHP1-), 5 minutes (p<0.05; Jurkat, J.PTPN22-, J.SHP1-) and 10 minutes (p<0.05; Jurkat) post stimulation, and this induction was ablated entirely in all cell lines after U0126 treatment (Figure 2H, Supporting Figure 7). Lastly, the activation site PLCγ1^Y783^ was significantly more responsive in non-U0126-treated J.TCPTP- cells at 2 minutes after TCR stimulation compared to all other cell lines (p<0.05). J.TCPTP- cells treated with U0126 showed significant phosphorylation of PLCγ1^Y783^ at all time points, whereas Jurkat and J.SHP1- only showed PLCγ1^Y783^ induction after 5- and 10-minutes, and J.PTPN22- showed no significant PLCγ1^Y783^ response (Figure 2I, Supporting Figure 8). Taken together, our selected clones J.TCPTP- (T2 1C10), J.PTPN22- (P1 1C1), and J.SHP1- (S1 2C4) are suitable for evaluating the regulatory role of TCPTP, PTPN22, and SHP1 in the context of pY homeostasis and TCR signalling.

**Figure 2:**
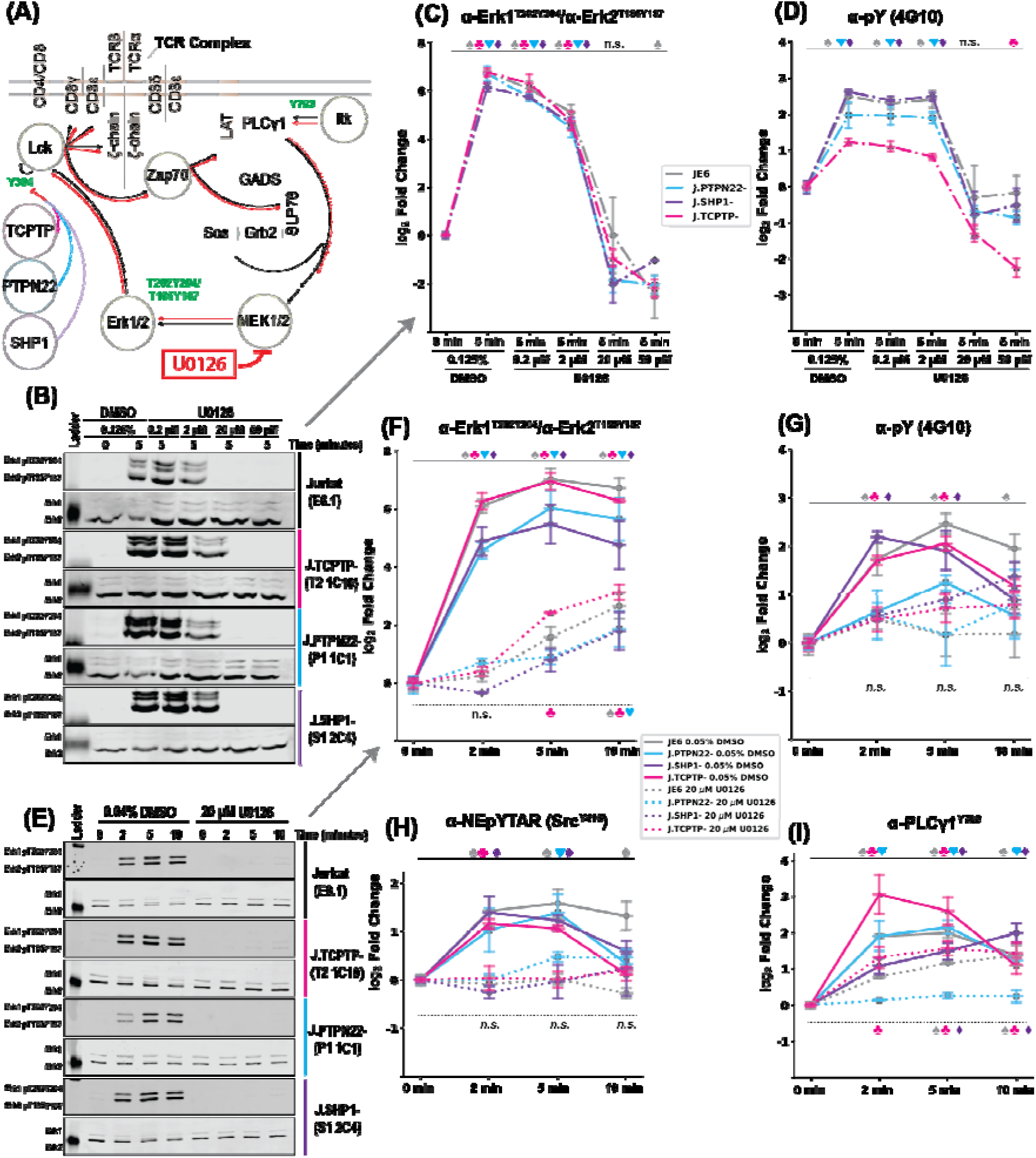
Knockout of *PTPN2* (TCPTP), *PTPN22* (PTPN22), or *PTPN6* (SHP1) retain TCR signalling and responsiveness to U0126 treatment. (A) Truncated schematic of TCR signalling, with black lines/arrows showing forward signalling and red lines/arrows showing the effect of U0126 treatment to reduce flux through the TCR signalling pathway. (B) Representative Western blots targeting Erk1^T202Y204^/Erk2^T185Y187^ during a U0126 titration. Quantification of U0126 titration Western blots targeting (C) Erk1^T202Y204^/Erk2^T185Y187^ and (D) pY. (E) Representative Western blots targeting Erk1^T202Y204^/Erk2^T185Y187^ during a TCR stimulation time course in the presence or absence of U0126. Quantification of time course Western blots targeting (F) Erk1^T202Y204^/Erk2^T185Y187^, (G) pY, (H) Src family kinase activation site, sequence NEpYTAR (G) PLCγ1^Y783^. For all Western blots, statistical significance between 0-minute normalised, log2 transformed abundances is determined by Fisher’s LSD with the Holm Sidak correction. ♠♣♥♦ indicate p<0.05 when comparing the given time point to the 0-minute sample for Jurkat, J.TCPTP-, J.PTPN22-, and J.SHP1-, respectively. Upper symbols are for 0.04% DMSO treatment, and lower symbols are for U0126 treatment time courses. ‘n.s’ indicates no statistical significance in that comparison for any cell line.

### TCPTP, PTPN22, and SHP1 play different regulatory roles on global pY abundance

Since PTPs are responsible for tyrosine dephosphorylation, which can activate or inactivate target proteins, we used the Src SH2 superbinder (sSH2) pY enrichment strategy^52,53^ to evaluate global levels of site-specific tyrosine phosphorylation in Jurkat, J.TCPTP-, J.PTPN22-, and J.SHP1- in the basal state and an antibody-stimulated state. PCA clustering showed cell line clustering and, to a lesser extent, clustering of the stimulated and unstimulated states for each cell line (Figure 3A). In the basal state, we found that few significantly changing pY sites were uniquely decreased in J.TCPTP- (15 sites) or J.PTPN22- (13 sites), and J.SHP1- showed a marked reduction in pY site abundance overall (123 sites) compared to WT Jurkats. In contrast, J.PTPN22- showed 260 unique pY sites significantly elevated, with few unique sites upregulated in J.TCPTP- (21) and J.SHP1- (25) (Figure 3B, 3C, 3D-E top row). In the stimulated state, 106, 51, and 3 pY sites were uniquely downregulated, and 52, 19, and 256 pY sites were uniquely upregulated in J.SHP1-, J.TCPTP-, and J.PTPN22-, respectively (Supporting Figure 9A-B, Figure 3D-E bottom row). PTPN22 KO significantly upregulated many canonical T cell signalling protein pY sites in both the basal and stimulated states, including ITAMs on TCR subunits (CD3δ^Y160^, CD3ε^Y188^, CD3γ^Y160^, TCRζ^Y142^, and TCRζ^Y153^), Lck^Y192^, activating sites on Zap70 (Zap70^Y492^, Zap70^Y493^, Zap70^Y492Y493^), the activating site Itk^Y512^, docking site LAT^Y220^ and GADS^Y45^.

**Figure 3:**
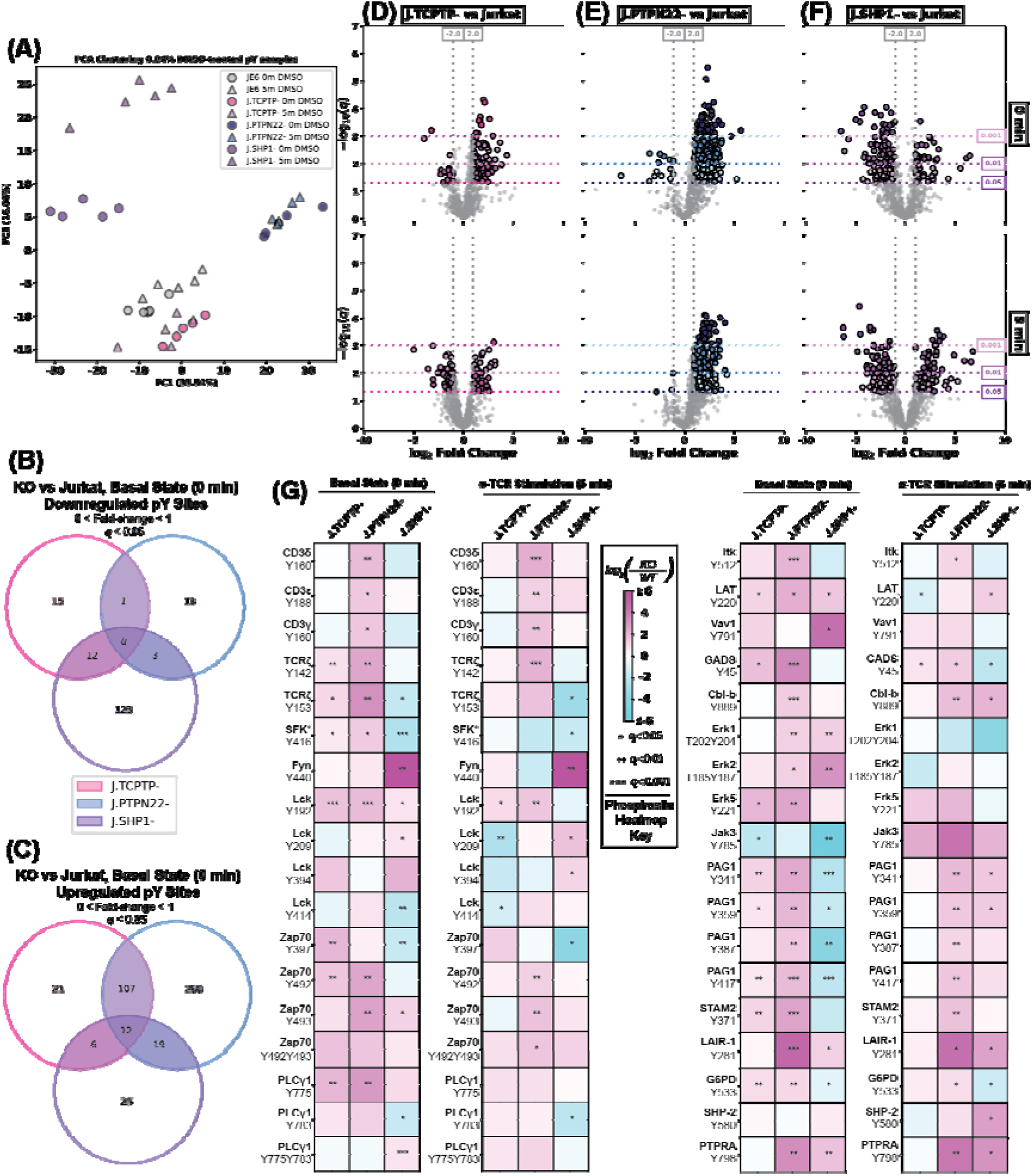
TCPTP, PTPN22, and SHP1 differentially maintain global pY abundance in Jurkat T cells. (A) PCA clustering on pY proteomics data. Venn diagram showing the overlap between (B) downregulated and (C) upregulated pY sites in J.TCPTP-, J.PTPN22-, and J.SHP1- in the basal state samples. (D-E) Volcano plots comparing unique pY sites in J.TCPTP-, J.PTPN22-, and J.SHP1-, respectively, to Jurkats in the basal state (0 minute - top row) and stimulated state (5 minute - bottom row). (G) Heatmaps showing individual pY sites compared with Jurkats for each phosphatase KO.

TCPTP KO displayed a milder yet similar phenotype as PTPN22, with specific upregulation of TCRζ, Zap70, and PLCγ1 phosphorylation sites that are no longer significantly upregulated after TCR stimulation. Interestingly, SHP1 KO led to a significant reduction in some canonical TCR phosphorylation sites, including TCR ζ^Y153^, the Src family activation site (peptide sequence LIEDNEpYTAR, SFK^Y416^), and PLCγ1^Y783^, while elevating other sites, including Erk1^T202Y204^/Erk2^T185Y187^, Vav1^Y791^, and Fyn^Y440^. After stimulation, most pY sites on PAG1—a negative-feedback adaptor that, once phosphorylated, recruits the Src-family-kinase inhibitor Csk—switched from significantly down-regulated to up-regulated in SHP1-deficient cells but not in the PTPN22- or TCPTP-knockouts (Figure 3G). This rise in PAG1 phosphorylation would be expected to bring Csk to the membrane, promote phosphorylation of Lck Tyr^505^, and thereby lock Lck in its inactive, closed conformation, providing an additional brake on TCR signalling uniquely evident when SHP1 is absent (Figure 3G). Using post-translational modification signature enrichment analysis (PTM-SEA), an algorithm that determines whether specific signatures are overrepresented or underrepresented in PTM-based proteomics data^54^, Fyn substrate signatures were elevated by TCPTP and PTPN22 KO, T cell receptor signatures were elevated after stimulation in J.TCPTP-, and □-CD3 signatures were elevated in all KOs after stimulation (Supporting Figure 9C-F). Together, our data suggest that TCPTP, PTPN22, and SHP1 each regulate unique aspects of T cell pY homeostasis, with differential influence on TCR signalling pY sites, contrary to the view that they are essentially redundant.

### SHP1 and PTPN22 play opposing roles in pY induction after TCR stimulation

Signalling from the TCR, whether through soluble antibody stimulation, soluble pMHC conjugation, or cell-based TCR-pMHC ligation, consists of characteristic pY site induction across many TCR signalling proteins, which dampen over time (> 10 minutes after ligation) due to engagement of negative feedback pathways^1^. After stimulation with □-Jurkat TCR□ (clone C305), we observed very few significantly downregulated pY sites in all cell lines, with J.TCPTP- having the most uniquely downregulated pY sites (38; Figure 4A). In contrast, J.SHP1- had a robust response to stimulation, with 121 pY sites significantly upregulated, compared with 12 and 6 pY sites unique to J.PTPN22- and J.TCPTP-, respectively (Figure 4B). Stimulation of cells led to a massive wave of pY induction, far greater than Jurkats or J.TCPTP-, while induction and significance of pY sites in J.PTPN22- was globally muted (Figures 4C-F), suggesting that SHP1 plays a uniquely inhibitory role in TCR signalling. We observed characteristic induction patterns for ITAMs on CD3δ/ε/γ and TCRζ, the activation site Zap70^Y492Y493^, an inhibitory site PLCγ1^Y771^, an activating site PLCγ1^Y775^, and GADS^Y45^, a site necessary for signal integration between the TCR and CD28, with SHP1-deficient cells showing the most potent induction. In J.SHP1- cells, PAG1 became uniquely responsive to TCR signalling across multiple pY sites, suggesting that SHP1 plays a unique role in preventing stimulation-dependent negative feedback to the TCR (Figure 4G). PTM-SEA revealed that, after receptor stimulation, Fyn, Lck, and Zap70 substrate signatures were significantly elevated in all cell lines after stimulation, with J.TCPTP- having the smallest enrichment for Fyn signatures (Figure 4H). T cell receptor signatures were significantly enriched in Jurkats, J.TCPTP-, and J.PTPN22- but not in J.SHP1- after receptor stimulation (Figure 4I), suggesting that loss of SHP1 increases the TCR responsiveness of typically non-TCR responsive pY sites. Finally, small molecule perturbation signatures consistent with activating agents, including □-CD3, were significantly elevated after stimulation in all cells, whereas inhibitor signatures, including U0126, were significantly decreased (Figure 4J).

**Figure 4:**
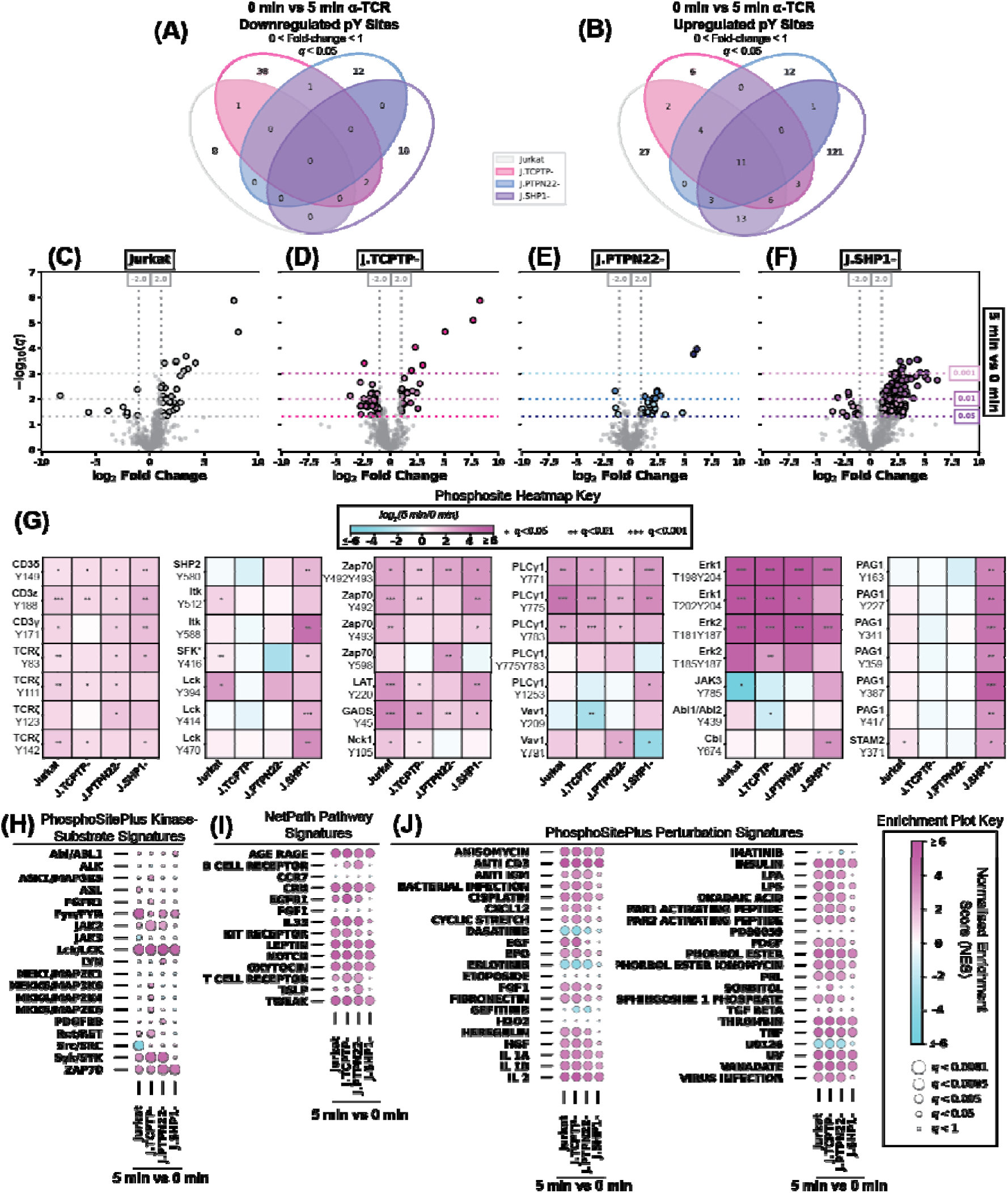
PTPN22 knockout mutes TCR signalling, while SHP1 knockout enhances TCR signalling. Venn diagram showing the overlap between (A) downregulated and (B) upregulated pY sites in Jurkat, J.TCPTP-, J.PTPN22-, and J.SHP1- after TCR stimulation. (C-F) Volcano plots comparing unique pY abundance in TCR stimulated (5 minute) and unstimulated samples in Jurkat, J.TCPTP-, J.PTPN22, and J.SHP1-. (G) Heatmaps showing fold-change and significance of individual pY sites in the TCR signalling pathway. (H-J) Bubbleplots showing post-translational modification signature enrichment analysis of PhosphoSitePlus Kinase Signatures, NetPath Pathway Signatures, and PhosphoSitePlus Perturbation Signatures, respectively, for comparisons between 5 minute and 0 minute samples.

Overall, these results suggest that genomic KO of TCPTP, PTPN22, or SHP1 does not prevent the forward progression of TCR signalling. However, PTPN22 KO slightly impairs TCR stimulation responsiveness, and SHP1 KO enhances it, consistent with the preliminary Western data (Figure 2).

### TCPTP knockout elevates pY abundance and decreases U0126 efficacy during TCR stimulation

U0126 treatment is known to interfere with TCR signalling due to positive feedback from Erk1/2. To evaluate the potential role of TCPTP, PTPN22, and SHP1 in Erk1/2 positive feedback, we treated Jurkats, J.TCPTP-, J.PTPN22-, and J.SHP1- with 20 µM U0126 before TCR stimulation with □-Jurkat TCR□ (clone C305) and sSH2 pY enrichment proteomics, as previously described^52,53^. Treatment with U0126 reduced the distinction in PCA clustering between basal state and stimulated samples while generally maintaining cell line clustering (Supporting Figure 10A). Compared to Jurkats, U0126-treated J.TCPTP- and J.PTPN22- had elevated pY levels in the basal and stimulated states, whereas J.SHP1- had reduced pY abundance in both (Supporting Figure 10B-D). The number of pY sites observed as significantly downregulated or significantly upregulated in the knockouts was similar to those observed without U0126 (Figures 3B-C, Supporting Figures 9A-B, 10A-D), which included many pY sites along the TCR signalling pathway (Supporting Figure 10E). During U0126 treatment, □-Jurkat TCR□ (clone C305) induced mild TCR stimulation in line with our previous work^21^, and PTPN22 deletion did not affect the pY induction. J.SHP1- cells showed no engagement along the TCR signalling pathway (Figure 5E-G), suggesting that the decrease of global phosphorylation due to SHP1 and U0126 treatment is muted after 5 minutes of stimulation compared to in the absence of U0126 (Figure 4). Meanwhile, □-Jurkat TCR□ (clone C305) stimulation of J.TCPTP- showed few significantly changing pY sites, but higher fold changes than any other cell line (Figures 5A-F). Many of the significantly changing pY sites were on T cell signalling proteins, including Zap70^Y492Y493^, GADS^Y45^, PLCγ1^Y783,^ and Erk1^T202Y204^. We observed enrichment of Fyn, Lck, and Zap70 kinase substrate signatures in all but J.SHP1- after stimulation (Figure 5G-H). We observed enrichment of various NetPath Pathway Signatures in J.TCPTP- and J.PTPN22-, however, TCR pathway signatures were downregulated in Jurkats and not enriched in J.TCPTP-, J.PTPN22-, or J.SHP1- after TCR stimulation, despite significant upregulation of □-CD3 signatures in J.TCPTP- and J.PTPN22- (Figure 5I-J). Together, our data show that even with the protective effect of TCPTP and PTPN22 knockouts on pY abundance, the loss of TCPTP is the most influential concerning U0126 treatment.

**Figure 5:**
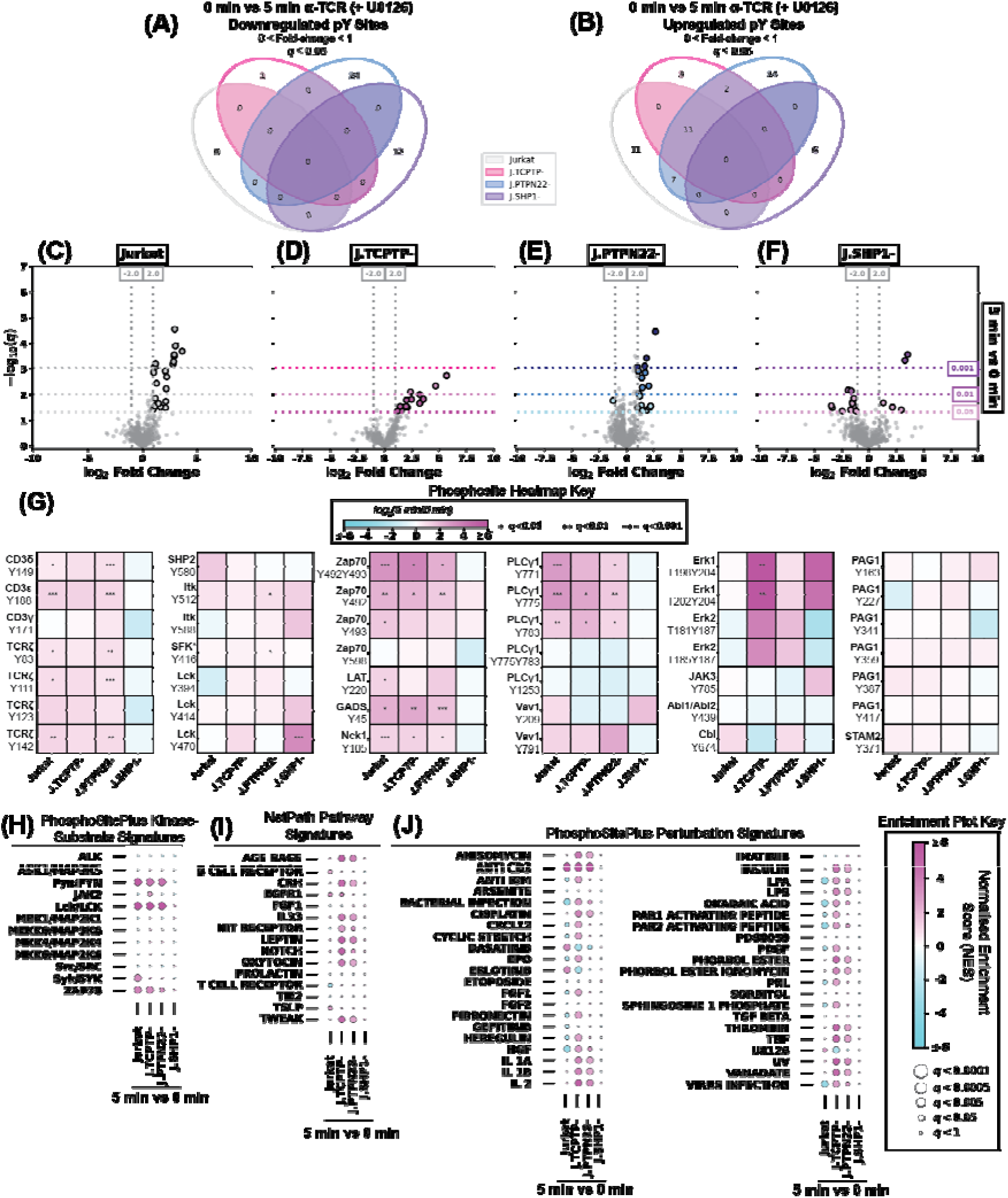
TCPTP recovers the highest fold-change induction during TCR stimulation with U0126 treatment. Venn diagram showing the overlap between (A) downregulated and (B) upregulated pY sites in Jurkat, J.TCPTP-, J.PTPN22-, and J.SHP1- after TCR stimulation during 20 μM U0126 treatment. (C-F) Volcano plots comparing unique pY abundance in TCR stimulated (5 minute) and unstimulated samples in Jurkat, J.TCPTP-, J.PTPN22, and J.SHP1- during 20 μM U0126 treatment. (G) Heatmaps showing fold-change and significance of individual pY sites in the TCR signalling pathway during 20 μM U0126 treatment. (H-J) Bubbleplots showing PTM-SEA of PhosphoSitePlus Kinase Signatures, NetPath Pathway Signatures, and PhosphoSitePlus Perturbation Signatures, respectively, for comparisons between 5 minute and 0 minute samples during 20 μM U0126 treatment.

### TCPTP knockout protects against U0126-dependent pY depletion

Our laboratory previously reported that MEK1/2 inhibition with U0126 leads to global depletion of tyrosine phosphorylation, a primary phenotype of Erk1/2 positive feedback in T cells^21^. We found the same results here; treatment with U0126 decreased pY abundance in Jurkat T cells in both the basal (140/8 down/up-regulated pY sites) and stimulated (200/3 down/up-regulated pY sites) states (Figure 6A, Supporting Figures 11A-C). The severe loss of pY abundance in Jurkats was phenocopied in J.PTPN22- and J.SHP1-, with few upregulated sites in the basal or stimulated states (Figures 6C-D, Supporting Figures 11A-C). Treatment of J.TCPTP- with U0126 led to a modest loss of pY abundance in the basal state (87/28 down/up-regulated pY sites) that was almost completely neutralised by TCR stimulation (14/17 down/up-regulated pY sites), suggesting that TCPTP loss protects against U0126-induced pY loss on a global level (Figures 6B, 5E, Supporting Figures 11A-C). In the basal state, U0126 treatment significantly lowered pY abundance on ITAMs (CD3δ^Y160^, CD3□^Y199^, TCRζ^Y142^), Erk1^T202Y204^, Zap70^Y492^, and other TCR signalling proteins. In general, pY sites on TCR signalling proteins were the most impacted in J.PTPN22-, with the additional downregulation of ITK^Y512^, GADS^Y45^, PLCγ1^Y775^, STAM2^Y371^, PAG1^Y163^, and GIT1^Y545^ (Figure 6F). In the stimulated state, the poor responsiveness to □-Jurkat TCR□ (clone C305) exacerbated the pY abundance loss, with the exception of J.TCPTP-. We observed upregulation of non-canonical pY sites on Lck (Lck^Y209^, Lck^Y414^), Itk (Itk^Y273^), and Zap70 (Zap70^Y248^), while observing very few pY sites on TCR signalling proteins with significant downregulation after U0126 treatment (Figure 6F). Using PTM-SEA, we found that U0126 treatment uniformly downregulated Fyn substrates with minimal effects on Lck substrates, and we observed no significant enrichment of Zap70 substrates (Figure 6G). As expected, U0126 treatment uniformly downregulated NetPath Pathway signatures, with significant downregulation of pY sites associated with TCR signalling in the basal and stimulated state of J.TCPTP- (Figure 5H). Finally, U0126 treatment led to significant downregulation of □-CD3 signatures across all cell lines and states and upregulation of U0126 signatures in the basal state and stimulated state of J.TCPTP- (Figure 6I). Together, these data suggest the existence of a pathway involving feedback from Erk1/2 through TCPTP that maintains pY balance in Jurkat T cells.

**Figure 6:**
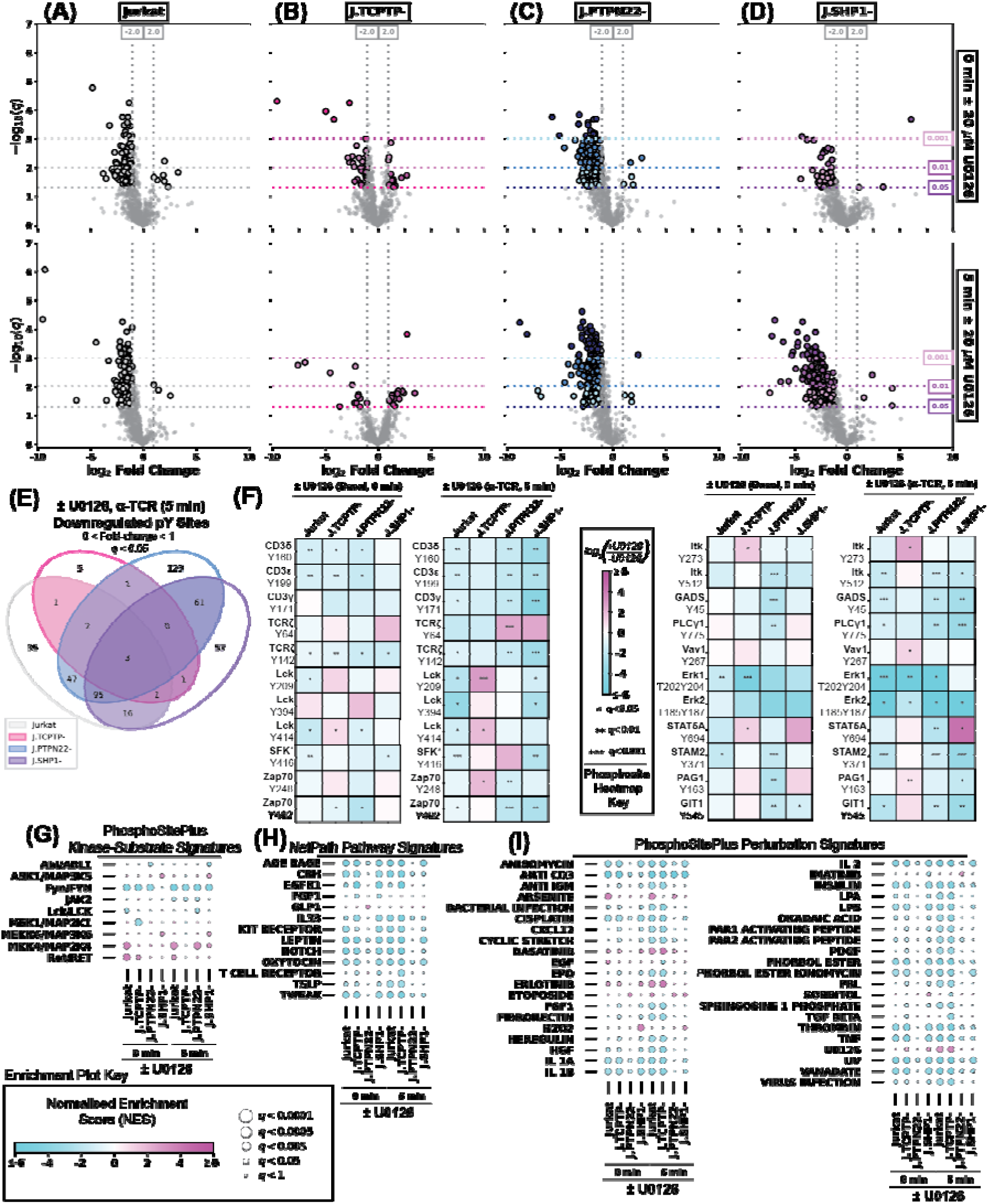
TCPTP plays a protective effect on pY abundance in the TCR-stimulated state during U0126 treatment. (A-D) Volcano plots comparing unique pY sites sequenced in Jurkat, J.TCPTP-, J.PTPN22-, and J.SHP1-, respectively, treated with 0.04% DMSO or 20 μM U0126. (E) Venn diagram showing the overlap between downregulated pY sites in Jurkat, J.TCPTP-, J.PTPN22-, and J.SHP1- in the 5 minute stimulation condition with or without 20 μM U0126 treatment. (F) Heatmaps showing fold-change and significance of individual pY sites in the TCR signalling pathway with or without 20 μM U0126 treatment. (G-I) Bubbleplots showing PTM-SEA of PhosphoSitePlus Kinase Signatures, NetPath Pathway Signatures, and PhosphoSitePlus Perturbation Signatures, respectively, for basal state and stimulated state comparisons with or without 20 μM U0126 treatment.

## Discussion

Regulation of cellular signalling pathways is necessary for effective and timely responses to external factors, including small molecule perturbation, alterations in nutrients, or influx of particular ligands. In the context of immune cell signalling, alterations in the kinetics, overall duration, or magnitude of signalling pathways cause autoimmune disorders^2,39^. Forward propagation of T cell receptor signalling, in which a pMHC complex on an antigen-presenting cell ligates a TCR for sufficiently long, is a well-studied process that requires the coordinated activation and deactivation of several enzymes, namely tyrosine kinases^2^. In particular, regulation of the initiating kinase, Lck, has been the subject of intense research and debate; five phosphatases (CD45 on Lck^Y505^, CD45/TCPTP/PTPN22/SHP1/JKAP on Lck^Y394^) and three kinases (Csk on Lck^Y505^, Lck on Lck^Y394^, Itk on Lck^Y192^) are known to act on Lck and control the forward progression of TCR signalling^3,5,7,8,10,12–16,18,51,55–58^. CD45 and Csk compete for Lck deactivation, as Lck^Y505^ locks Lck in an inactive conformation, preventing autophosphorylation of Lck^Y394^ and subsequent activation of Lck. The four phosphatases TCPTP, PTPN22, SHP1, and JKAP have all been shown to dephosphorylate Lck^Y394^ *in vitro*. Conflicting evidence exists for the ‘primary’ phosphatase responsible for Lck^Y394^ dephosphorylation^13–15,45,55^. Complications arise from overwhelming evidence for a positive feedback loop involving the kinases Erk1/2 and Lck that maintains pY levels in T cells, which is thought to involve Lck^S59^ phosphorylation by Erk1/2 that disrupts SHP1 binding despite contradictory evidence^21,23,45^. Fewer studies have evaluated the regulation of the other TCR proximal kinases, Fyn, Zap70, and Itk/Tec, leading to ambiguities in the feedback mechanisms involved in accurate and timely TCR signalling that is crucial for an effective immune response. Our work shows that the phosphatases TCPTP, PTPN22, and SHP1 have distinct regulatory roles in the context of global pY maintenance and regulation of the T cell receptor.

T cell protein tyrosine phosphatase was one of the first characterised phosphatases in T cells and exists in 48 kDa nuclear- and 45 kDa cytoplasmic-lolising isoforms^59–61^. Early work described TCPTP-mediated Lck dephosphorylation^15^ and regulation of Janus kinase 1/3^62^, implicating TCPTP as a key negative regulator of T cell activation pathways^63^. While work investigating the specificity of TCPTP was performed early^64^, signalling studies of TCPTP in T cells have largely ceased despite the immunological relevance of TCPTP. Recent work evaluating TCPTP has focused on TCPTP structure^65–68^, disease-relevant mutations, and drugs^44,63,69–72^. Our work suggests that TCPTP is a modest negative regulator of T cell receptor signalling (Figure 3), and TCPTP loss allows for recovery of TCR induction through a PLCγ1-dependent pathway (Figure 2). Further, loss of TCPTP attenuates the pY loss associated with Erk deactivation (Figure 6) and increases fold-induction after TCR stimulation (Figure 5). Four non-canonical pY sites on key TCR signalling proteins - Lck^Y209^, Lck^Y414^, Zap70^Y248^, and Itk^Y273^ - are significantly elevated in the TCR stimulated state when J.TCPTP- are treated with U0126 (Figure 6F), providing possible phosphorylation sites regulated by TCPTP that are involved in Erk feedback. Notably, Y^273^ (Itk), Y^248^ (Zap70), and Y^209^ (Lck) lie adjacent to, or within, the SH2 domain of their respective kinases; analogous to the well-characterised Itk-dependent phosphorylation of Lck Y^192^, these modifications could modulate SH2-mediated docking specificity and rewire protein-interaction networks within the T-cell signalosome^73^. Thus, TCPTP may be involved in positive feedback mechanisms involving MEK1/2 and Erk1/2 to maintain pY homeostasis and TCR responsiveness in T cells (Figure 7).

**Figure 7:**
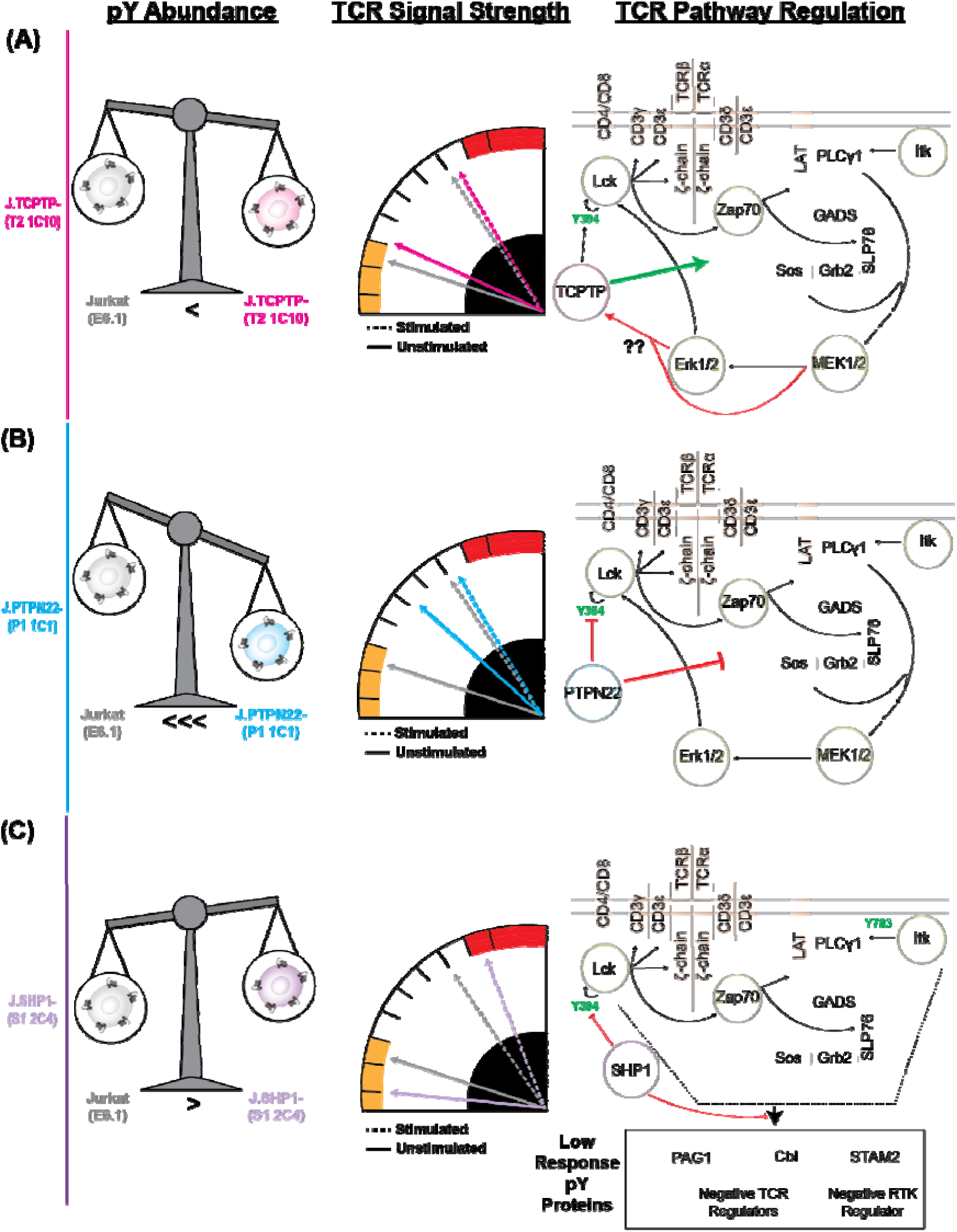
A model depicting the effects of (A) TCPTP, (B) PTPN22, and (C) SHP1 on global pY abundance, TCR stimulation strength, and TCR pathway regulation. Global pY abundance increases slightly upon deletion of *PTPN2*, largely upon deletion of *PTPN22*, and decreases slightly upon *PTPN6* deletion. J.TCPTP-cells respond similarly to Jurkats when stimulated with α-Jurkat TCRα, whereas J.PTPN22-cells have a muted response, and J.SHP1- cells have an elevated response. Loss of TCPTP ameliorates Erk positive feedback in the stimulated state, thus TCPTP is involved in a feedback pathway involving Erk1/2 and/or MEK1/2. Loss of PTPN22 elevates pY abundance globally and mutes TCR responsiveness, thus PTPN22 acts as a major negative regulator of the TCR signalling pathway. Loss of SHP1 slightly elevates TCR signalling pathway pY abundance while altering the regulation of negative TCR regulators and globally reducing pY abundance, thus SHP1 regulates other forms of negative feedback to the TCR.

Protein tyrosine phosphatase non-receptor type 22 (PTPN22; also lymphoid tyrosine phosphatase Lyp) was first isolated in 1993 by Cohen *et al*. and later identified to interact with Csk and dephosphorylate various TCR signalling proteins^16–18,29,74–76^, cementing the role of PTPN22 as a negative regulator of T cell receptor signalling. Mutations in PTPN22 were then increasingly documented in cases of autoimmune disorders^24,27,32,34,38,40,41^, and PTPN22 is being investigated as a druggable target for cancer immunotherapies^26,77^. Our work is well aligned with previous studies: PTPN22 knockout elevates TCR pY levels, mutes TCR responsiveness, and enhances the effects of MEK1/2 inhibitor (Figures 2-7). Interestingly, *PTPN22* deletion elevates the abundance of LAIR-1^Y281^, an ITIM site required for SHP1 binding^78,79^, indicating a potential method of phosphatase crosstalk in TCR regulation. However, our work demonstrates the broader, global effect of PTPN22 on resting pY levels in a TCR-independent context, suggesting that the role of PTPN22 may extend beyond just T cell signalling.

Src homology 2 domain-containing phosphatase 1 (SHP1) was first isolated in 1992^80^ and subsequently characterised as a negative regulator of various immune signalling pathways, including TCR signalling. In the TCR signalling context, SHP1 dephosphorylates Lck^Y394^ and can associate with unphosphorylated Lck^S59^ to downregulate TCR signalling activity^11,45,81–83^. SHP1 also binds directly to the immunoreceptor tyrosine-based inhibitory motifs (ITIMs) on PD-1 and CEACAM1^84,85^, bringing negative TCR signalling regulators to the active TCR signalosome and preventing exhaustion^86^. Our work agrees with previous studies that SHP1 acts as a negative regulator of TCR signalling, as *PTPN6* deletion results in upregulation of various TCR signalling phosphorylation sites Lck^Y192^, Lck^Y394^ (trending but n.s. in all knockouts), Zap70^Y493^, LAT^Y220^, PLCγ1^Y775Y783^, and SHP2^Y580^ in the stimulated and/or basal states (Figure 3G). In contrast, global pY abundance appears to drop without SHP1 (Figure 3), and more pY sites become strongly TCR responsive (Figure 4). Some pY proteins, including PAG1 and Cbl, are known negative regulators of TCR signalling^87,88^, suggesting that SHP1 may play a context-dependent role in suppressing negative regulators of the TCR. Although the role of SHP1 in T cell pY maintenance and TCR signalling is still evolving, SHP1 is implicated in many autoimmune disorders and cancers^25,31,36,89–92^, thus understanding the mechanisms of SHP1 regulation of T cell signalling is imperative.

Regulation of T cell activation is a critical process for maintaining a coordinated immune response. Loss-of-function mutations in the PTPs TCPTP, PTPN22, and SHP1 cause autoimmune disorders through immune system dysregulation, particularly in T cells and their responsiveness to antigens. The redundancy of these phosphatases, coupled with the small number of TCR-responsive pY sites, raises questions regarding the timing, specificity, and substrates of these proteins. Our work addresses the questions of timing and specificity: each phosphatase has a distinct influence on the phosphotyrosine proteome in T cells, and each controls a subpopulation of TCR-responsive pY sites. Further, we show that TCPTP has a protective effect on MEK1/2 inhibition, a role unique to TCPTP. To better understand the unique role each phosphatase plays in T cell biology, it is necessary to investigate the specific substrates for each phosphatase and their effects on those pathways. Because SHP1^93^, TCPTP^94^ and PTPN22^95,96^ are now being pursued as drug targets, our mechanistic insights have immediate translational relevance. Specifically, the finding that TCPTP loss attenuates the broad pY depletion normally caused by MEK1/2 inhibition suggests that cancers or immune cells with low TCPTP activity could display primary resistance to MAP□kinase–pathway inhibitors—yet might be rendered vulnerable by dual blockade of TCPTP and MEK/ERK. Conversely, the global pY elevation and muted TCR responsiveness we observe upon PTPN22 deletion nominate easily quantifiable phosphotyrosine signatures that could act as pharmacodynamic biomarkers for forthcoming PTPN22□directed therapies. Together, these observations underscore the value of biomarker□guided combination strategies that account for the distinctive feedback circuits controlled by each phosphatase.

## Methods

### Cell Culture and Generation of PTP Knockout Cells

Jurkat T cells (clone E6.1, ATCC #TIB-152) and all JE6 derivatives were maintained in RPMI 1640 with 10% FBS (PeakSerum #PS-FB3), 2 mM L-glutamine, 100 U/mL penicillin, 100 μg/mL streptomycin (PSQ; HyClone #SV30082.01), and 10 mM HEPES, pH 7.4, at 5% CO_2_ in a humidified incubator. Jurkat T cells with a *PTPN22* (J.PTPN22-), *PTPN2* (J.TCPTP-), or *PTPN6* (J.SHP1-) genomic deletion were generated by CRISPR Cas9 by using guide RNAs targeting *PTPN22* (Forward = CACCGTCTTGCTTTGGGCCTCATCC, Reverse = AAACGGATGAGGCCCAAAGCAAGAC), *PTPN2* (Forward = CACCGTTCGAACTCCCGCTCGATGG, Reverse = AAACCCATCGAGCGGGAGTTCGAAC), and *PTPN6* (Forward = GTGAGTTCTGGATCCGAATAT, Reverse = AAACATATTCGGATCCAGAACTCAC) . Guide RNAs were cloned into the pSpCas9(BB)-2A-GFP vector or the PX458 vector before transfection by electroporation (260 V, 1250 uF) and a recovery period of 48 hours in RPMI 1640 with 10% FBS and no antibiotics. Single cells were sorted for high CD3 expression, an integral component of the T cell receptor complex, before expansion of single-cell clones. Clones were screened for PTPN22, TCPTP, or SHP1 expression by Western blot before sequencing to confirm the disruption of the gene.

### Flow cytometry

To evaluate the expression of cell surface receptors in the PTP knockout Jurkats, we collected one million cells by centrifugation, resuspended in flow cytometry buffer (2% FBS in 1X DPBS), and then incubated in fluorescently labelled antibodies for 30 minutes on ice in the dark. After receptor staining, the cells were washed 3 times in 1 mL, resuspended in 0.1 mL of flow cytometry buffer, then fixed with 2% PFA in the dark for 15 minutes at room temperature. PFA was washed off the cells 3 times with 1 mL flow cytometry buffer, then resuspended in 0.1 mL flow cytometry buffer before analysis on a Cytek Aurora flow cytometer. The resulting flow cytometry data were analysed and plotted using Floreada. Antibodies: CD3ε PE (1:50, BD Pharmingen #555333), TCRα PE (1:50, BD Pharmingen #555548), CD45 FITC (1:50, eBioscience #11-0459-41).

### Inhibitor Treatment and TCR Stimulation

Inhibition of MEK1/2 was performed using U0126 as previously described^21^. Briefly, a stock of U0126 (Cell Signaling Technology cat #9903) was resuspended at a concentration of 50 mM in 100% DMSO under sterile conditions and diluted in unsupplemented RPMI 1640 at a concentration of 20 uM (0.04% DMSO) before use with cells. A control treatment of 0.04% DMSO without U0126 was included for all treatments. For phosphotyrosine mass spectrometry analysis, 1 × 10^8^ cells were used, and for small-scale experiments, 1 × 10^7^ cells were used. For mass spectrometry experiments, a minimum of four biological replicates were used per condition, and for small-scale experiments, a minimum of four biological replicates were used per condition. Cells were harvested by centrifugation before resuspension in U0126 RPMI or control RPMI at a concentration of 1 × 10^6^ cells per millilitre for 2.5 hours. After treatment, cells were washed in warm 1X DPBS twice before resuspension at 2 × 10^8^ cells per millilitre and a 30-minute rest period. T cell receptor stimulation was initiated as previously described by a 1-to-1 addition of a-Jurkat TCRa (Clone C305, a gift from Arthur Weiss) for a final concentration of 1 ug/mL and allowed to proceed for 5 minutes before lysis. Basal state samples were prepared by the addition of urea lysis buffer, vigorous vortexing, and then addition of a-TCRa antibody solution to the lysed samples to maintain cell equivalence amongst samples.

### Lysis, Reduction, Alkylation, and Digestion for Proteomics

All samples were lysed using urea lysis buffer (8 M urea, 20 mM HEPES pH 7.4, 1 mM sodium orthovanadate, 2.5 mM sodium pyrophosphate, 1 mM µ’-glycerophosphate) and vigorous vortexing. Samples were incubated on ice for at least 15 minutes before sonication at 70% amplitude for 30 seconds, twice. Samples prepared for Western blot analysis were prepared by a 1-to-1 dilution in 2X Laemmli sample buffer (2% SDS, 20% Glycerol, 0.01% bromophenol blue, 0.125 M Tris-HCl, 5% µ’-mercaptoethanol) followed by 5 minutes at 95 C. Protein concentrations were determined using the Pierce BCA Assay kit (Thermo #23225) using the manufacturer’s instructions. The total protein across all samples for phosphotyrosine proteomics was adjusted to 5 mg per replicate, with a total of 5 biological replicates per condition. For global phosphoenrichment, samples were normalised after data acquisition to whole-cell lysate distributions. Urea lysates were reduced using 10 mM dithiothreitol for 30 minutes at 37 C and subsequently alkylated using 10 mM iodoacetamide for 30 minutes at room temperature. Lastly, samples were digested using sequencing grade modified trypsin (Promega #V5113) in a 1:50 (w/w) trypsin to protein ratio overnight at 37 C. Tryptic peptides were acidified at a final percentage of 1% (v/v) trifluoroacetic acid (TFA), centrifuged at 1800Xg at room temperature for 5 minutes to collect debris, then loaded onto Sep-Pak C18 vacuum cartridges (Waters #WAT020515) for desalting^97^. Samples were desalted in parallel with 12 mL of 0.1% TFA, eluted using 7 mL of 0.1% TFA in 40% acetonitrile, then diluted 1:1 in 0.1% TFA to reduce the acetonitrile concentration to 20%, frozen at -80 C overnight, and lyophilised for 48 hours before phosphoenrichment.

### Superbinder SH2 pY Enrichment

Selective enrichment of pY-containing peptides was performed using the superbinder SH2 (sSH2) domain as described previously^52,53^. Briefly, recombinant sSH2 was purified and conjugated to cyanogen bromide-activated sepharose beads (GE Healthcare, Wauwatosa, Wisconsin) for pY peptide capture. Desalted samples were resuspended in 3 mL IAP buffer (10 mM sodium phosphate monobasic monohydrate, 50 mM sodium chloride, 500 mM MOPS, pH 7.2) before a 2-hour incubation with about 1 mg of sSH2-containing beads at 4 C on an end over end rotator. After incubation, the beads were washed with 1 mL of ice-cold IAP buffer thrice, and then 1 mL of ice-cold HPLC-grade water thrice. Peptides were eluted from the sSH2 beads with 0.15% TFA on a 1,150 rpm thermomixer for 10 minutes at room temperature twice before desalting with Pierce C18 tips (Thermo #87784) according to the manufacturer’s instructions. Peptides were then dried by speed-vac and stored at -80 C until LC-MS/MS analysis.

### Liquid Chromatography-Tandem Mass Spectrometry

Phosphotyrosine-enriched peptides were separated on a lab-packed C18 column (150 mm by 75 μm) with XSelect CSH C18 2 μm resin using an Ultimate 3000 RSLCnano system. Peptides were eluted using a 90-minute method starting with a Buffer A (0.1% formic acid in HPLC water) to Buffer B (0.1 % formic acid in acetonitrile) ratio of 98:2 and progressing to a final ratio of 70:30 in 69 min, with a 90% buffer B spike at the end of the gradient. Peptides were analysed on an Orbitrap Exploris 480 mass spectrometer in positive ion mode with a spray voltage of 2.2kV, funnel RF level of 40, and heated capillary temperature of 275 C. Precursor mass range was set to 375-1500 m/z with a resolution of 120,000, normalised AGC target of 300%, IT of 50 ms in profile mode. Fragmentation spectra were collected using a cycle time DDA mode of 2.5 seconds with a dynamic exclusion time of 20 seconds, and an intensity threshold of 50,000. Monoisotopic precursor ions with charge states between +2 and +6 were fragmented using higher-energy collision-induced dissociation (HCD) using a normalised collision energy of 30%. Fragmentation spectra were acquired using a 0.7 m/z isolation window at a resolution of 15,000 and a normalised AGC target of 100% in centroid mode with an injection time of 100 ms.

Whole-cell digest peptides were separated on an Aurora C18 column (250 mm by 75 μm; IonOpticks #AUR3-25075C18) with 1.7 μm C18 using an Ultimate 3000 nanoRSLC. Peptides were eluted using a 120-minute method starting with a Buffer A (0.1% formic acid in HPLC water) to Buffer B (0.1% formic acid in acetonitrile) ratio of 98:2 and progressing to a final ratio of 70:30 in 69 min, with a 90% buffer B spike at the end of the gradient followed by a short wash gradient from 2% to 99%. Peptides were analysed on a QExactive mass spectrometer in positive ion mode with a spray voltage of 2.1kV, funnel RF level of 40, and heated capillary temperature of 275 C. Precursor mass range was set to 400-1800 m/z with a resolution of 70,000, AGC target of 3e6, IT of 200 ms in centroid mode.

Fragmentation spectra were collected using a Top9 DDA method with a dynamic exclusion time of 30 seconds, and an intensity threshold of 1e3. Monoisotopic precursor ions with charge states between +2 and +5 were fragmented using higher-energy collision-induced dissociation (HCD) using a normalised collision energy of 30. Fragmentation spectra were acquired using a 2.5 m/z isolation window at a resolution of 17,500 and an AGC target of 2e4 in centroid mode with a maximum injection time of 200 ms.

### Database Searching and Relative Phosphopeptide Abundance

For sSH2 enriched samples, Thermo .RAW files were analysed using the high throughput autonomous proteomics pipeline (HTAPP) and PeptideDepot, custom in-house software for the automation of PTM proteomics analysis^98,99^. MS2 spectra were searched using Mascot against the UniProt complete proteome dataset (93,802 nonredundant forward sequences, downloaded August 2019) to 0.1% FDR using a reverse decoy database. The following parameters were used for the search: trypsin enzyme cleavage (up to 2 missed cleavages), 7 ppm precursor mass tolerance, 20 mmu fragment ion mass tolerance, variable modifications for phosphorylation (S/T/Y; +79.9963 Da), oxidation (M; +15.9949 Da), and static modification of carbamamidomethylation (C; +57.0215 Da). Retention time alignment and integration of selected ion chromatograms for each peptide spectrum match were used to determine relative peptide abundance as previously described^100,101^.

Raw files were evaluated for whole-cell digest samples using MaxQuant (Version 2.1.3) with the integrated peptide search engine Andromeda, as described previously^102,103^. Briefly, MS2 spectra were searched against the human UniProt database (*Homo sapiens*, downloaded August 2019) containing 93,802 forward protein sequences. For peptide spectrum matches and proteins, the false discovery rate (FDR) upper threshold was set to 0.01% and was determined using a reverse sequence decoy database approach. The following modifications were set for database searching: carbamidomethylation (cysteine) was fixed, oxidation (methionine) was variable, acetylation (protein N-terminus) was variable, phosphorylation (serine, threonine, tyrosine) was variable. Trypsin specificity was used, and a maximum of two missed cleavages were tolerated. For the main search, the peptide tolerance was set to 7 ppm, whereas the MS/MS tolerance was set to 20 ppm. The search parameter files (mqpar.xml), and the search results Phospho (STY)Sites.txt, proteinGroups, and evidence files are available at the PRIDE partner repository under the dataset identifier PXD058624.

### Gel Electrophoresis and Western Blotting

Samples were separated on 10% Tris-Glycine polyacrylamide gels cast in-house using the BioRad Mini-Gel apparatus and transferred to a 0.45 um polyvinylidene difluoride Immobilin membrane (Millipore Sigma #IPFL00010) before blocking using Intercept TBS Blocking Buffer (LI-COR Biosciences #927-60003). Blocked membranes were incubated with a primary antibody solution containing 5% BSA in 1X Tris-buffered saline with Tween 20 (TBST) at 4 C overnight. After primary antibody incubation, membranes were washed three times with 1X TBST before incubation with a secondary antibody solution for 1 hour at room temperature. After secondary antibody incubation, the membranes were washed three times with 1X TBST and then washed twice with 1X TBS before visualisation using the Odyssey CLx and Odyssey M Imaging Systems (LI-COR Biosciences). Western blots were run in 2- replicate pairs for each group for a total of 4 replicates, and quantification was performed by adjusting the signal to the loading control, as previously described^104^, normalising to the 0- minute group, then combining replicate values between two blots before log_2_ transformation and statistical analysis. Due to changes in antibody lots and catalogue numbers for some targets (SFK^Y416^, Vav1), a direct comparison between cell lines was not performed, and all data presented are relative to their respective 0 minute/no treatment samples. Statistical significance between the mean log_2_ transformed, basal-state normalised fold changes was performed using a one-way ANOVA with Fisher’s LSD and the Holm Sidak FWER correction^105–108^.

Primary Antibodies: pY clone 4G10 (1:2000, Cell Signalling Technologies #96215), PLCγ1 (1:10000, Millipore Sigma #05-163), PLCγ1^Y783^ (1:2000, Cell Signalling Technologies #2821), Erk1/2 (Cell Signalling Technologies #9107), Erk^T202Y204^ (1:2000, Cell Signalling Technologies #9101), Lck (1:5000, Cell Signalling Technologies #2657S), Src Family Kinase Y416 (1:2000, Cell Signalling Technologies #59548S, 2101S), TCPTP (1:2000, Cell Signalling Technologies #58935S), PTPN22 (1:1000, Thermo #11783-1-AP), SHP1 (1:500, Santa Cruz Biotechnology #sc-287), and Vav1 (1:2000, Cell Signalling Technologies #4657S, 2505S) Secondary Antibodies: IRDye 680RD Goat anti-Mouse IgG (LI-COR #926-68070) IRDye 800CW Goat anti-Mouse IgG (LI-COR #926-33210), IRDye 680RD Donkey anti-Rabbit IgG (LI-COR #926-68073), and IRDye 800CW Donkey anti-Rabbit IgG (LI-COR #926-32213).

### Data Analysis & Code Availability

Post-search analysis of LC-MS/MS data was performed on Ubuntu 22.04 LTS on the Windows Subsystem for Linux (version 2) using Python (version 3.12.5) with the following dependencies: matplotlib (version 3.9.1), scipy (version 1.14.0), numpy (version 2.0.1), scikit-learn (version 1.5.1), pandas (version 2.2.2), MissingPy (modified for updates with scikit-learn), py-venn, and a custom ‘helpers’ module.

The “proteinGroups.txt” file from MaxQuant was used to analyse protein profiling data. The proteinGroups table was filtered to remove “Reverse” and “Potential contaminants” flagged rows, and proteins with >10% sequence coverage were kept for further analysis. Missing values were imputed using the Random Forest method using the MissingPy module^109,110^. Welch’s T tests^111^ with Benjamini & Hochberg’s FDR correction^112,113^ were performed to compare the mean log-transformed abundances of each cell line. For plotting, only rows containing >3 measured values were included. Replicate clustering and reproducibility were evaluated using principal component analysis and multiple linear regression after imputation. KEGG pathway enrichment was performed using ssGSEA2.0 with the KEGG Msig database^54^.

For pY data, the unique pY sites and their abundances were exported from PeptideDepot. Missing values were imputed as described above, and statistical significance was determined as described above, except comparisons were made as shown in Figures 3-6, Supporting Figure 10. Principal component analysis and linear regression were used to show replicate clustering and reproducibility, and ssGSEA2.0 was used to show pathway enrichment using the PTMsig database^54^.

## Acknowledgements

The authors would like to thank the Proteomics Core Facility at the University of Arkansas Medical School for collecting phosphotyrosine proteomics data, and the Proteomics Core Facility at Brown University for collecting whole cell digest proteomics data. We acknowledge the NIH grants R01AI083636, P01AI091580, P20GM121293, S10OD036295, and R24GM137786 for financial support for our research. We also thank Dr. Arthur Weiss and his laboratory for providing □-Jurkat TCR□ (clone C305). We wish Dr. Weiss a happy retirement and thank him for our fruitful collaborations over the years, especially all of the emails saying ‘Hi Art … Art’.

## Data and Code Availability

All mass spectrometry data have been uploaded to the ProteomeXchange Consortium via the PRIDE^114^ partner repository under the dataset identifier PXD058624. All code used for the analysis and presentation of our mass spectrometry data are publicly available on GitHub (https://github.com/Aurdeegz/Phosphatase-Project/) and are available upon request.

## Author Contributions

Conceptualisation: AC, ARS, MH

Methodology: AC, ARS, CH, NM, SPR

Investigation: AC, CH, AM, AW, AAG, XY, NH, RZP

Visualisation: AC

Funding acquisition: ARS Project administration: AC, ARS

Supervision: AC, ARS

Writing – original draft: AC

Writing – review & editing: AC, ARS, SPR

