## Supplementary Information for "The phosphatases TCPTP, PTPN22, and SHP1 play unique roles in T cell phosphotyrosine maintenance and feedback regulation of the TCR"

**Title:**

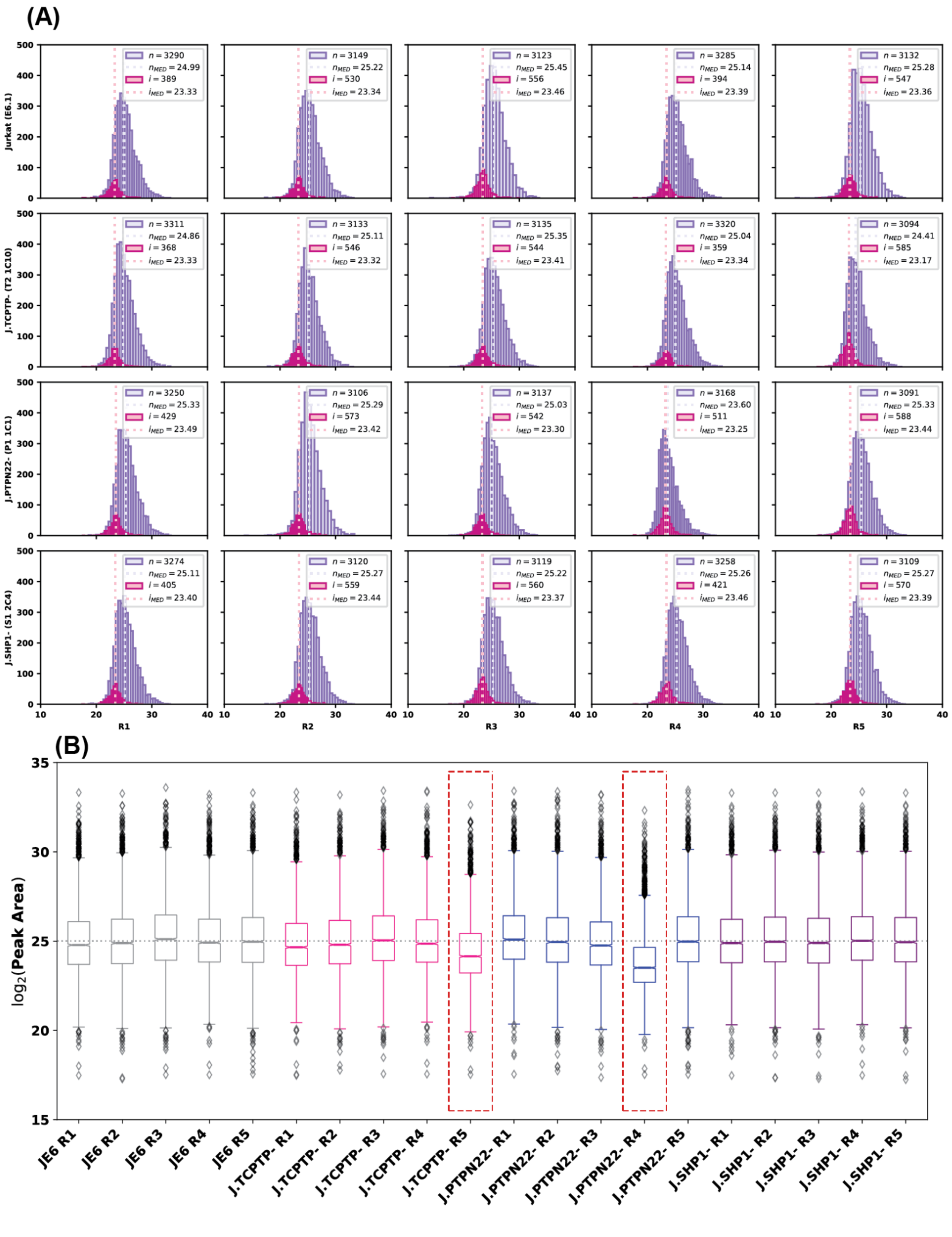

**Supporting Figure 1:** Quality control of Jurkat, J.TCPTP-, J.PTPN22-, and J.SHP1- protein profiling mass spectrometry data. (A) Histogram of log2 transformed abundances in purple with imputed distributions in pink. (B) Box and whisker plots showing the intensity distribution for each protein profiling sample. J.TCPTP- R5 and J.PTPN22- R4 are circled in red. Due to the notably lower protein abundance in these replicates, they were discarded from further analysis.

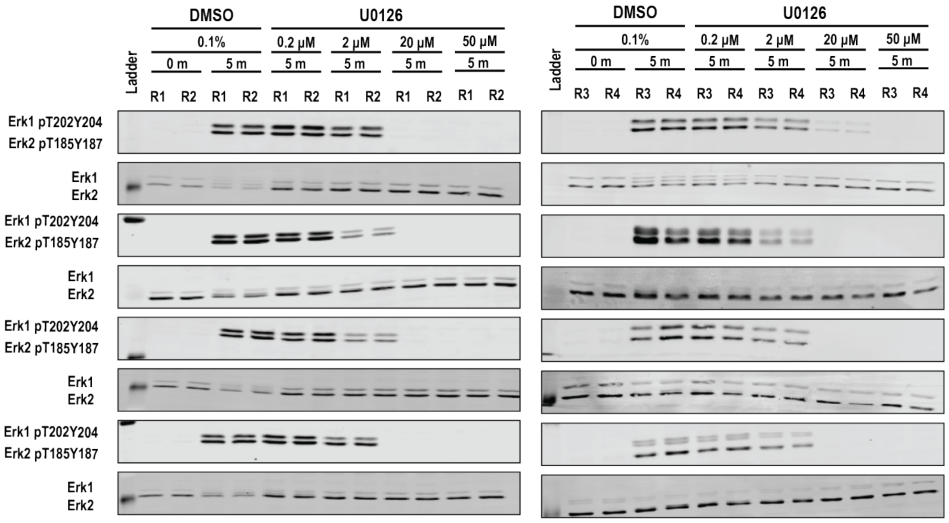


**Supporting Figure 2:** Western blots used for quantification of Erk1$^{T202Y204}$/Erk2$^{T185Y187}$ during a U0126 titration.

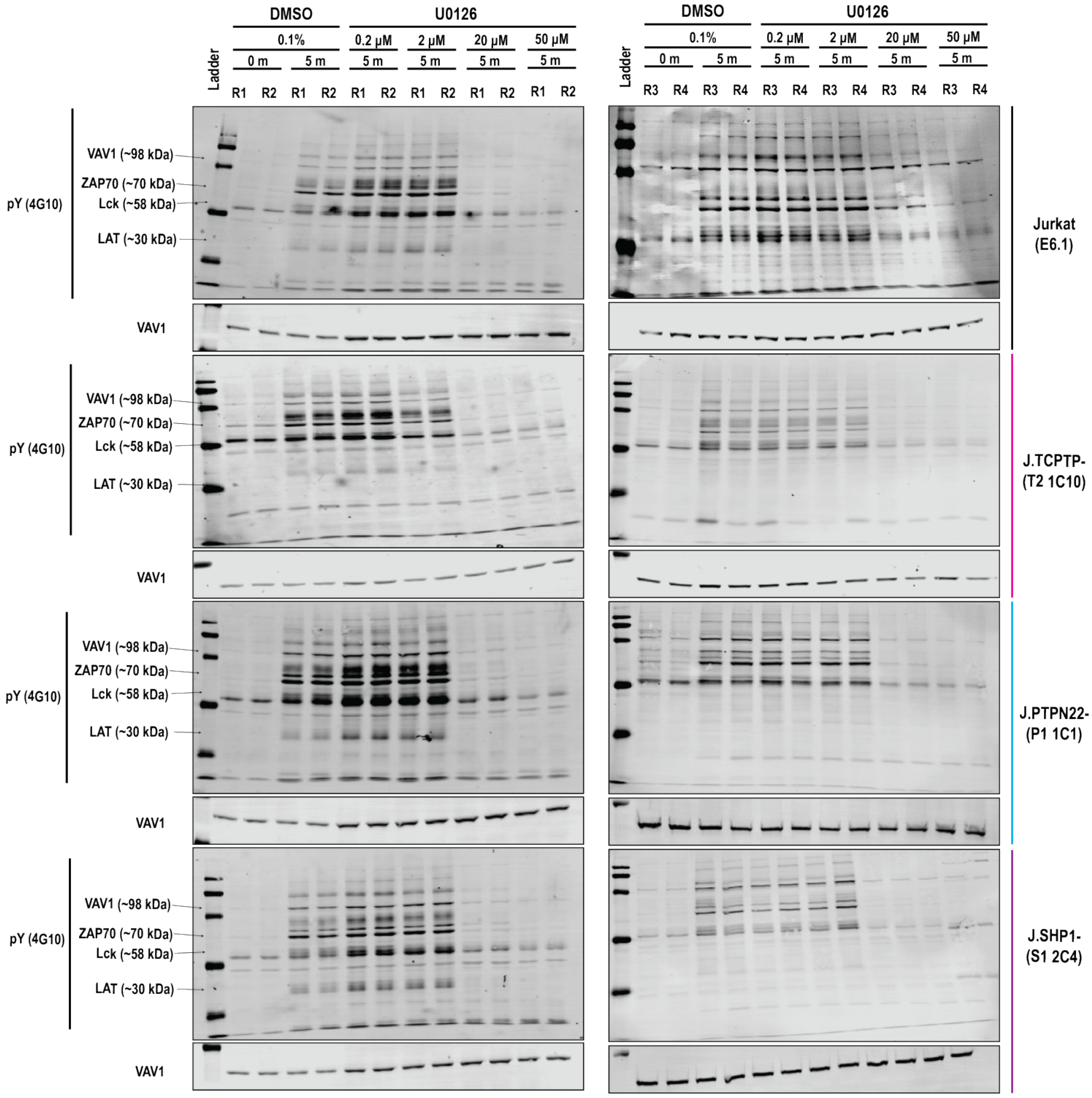


**Supporting Figure 3:** Western blots used for quantification of global pY abundance during a U0126 titration.

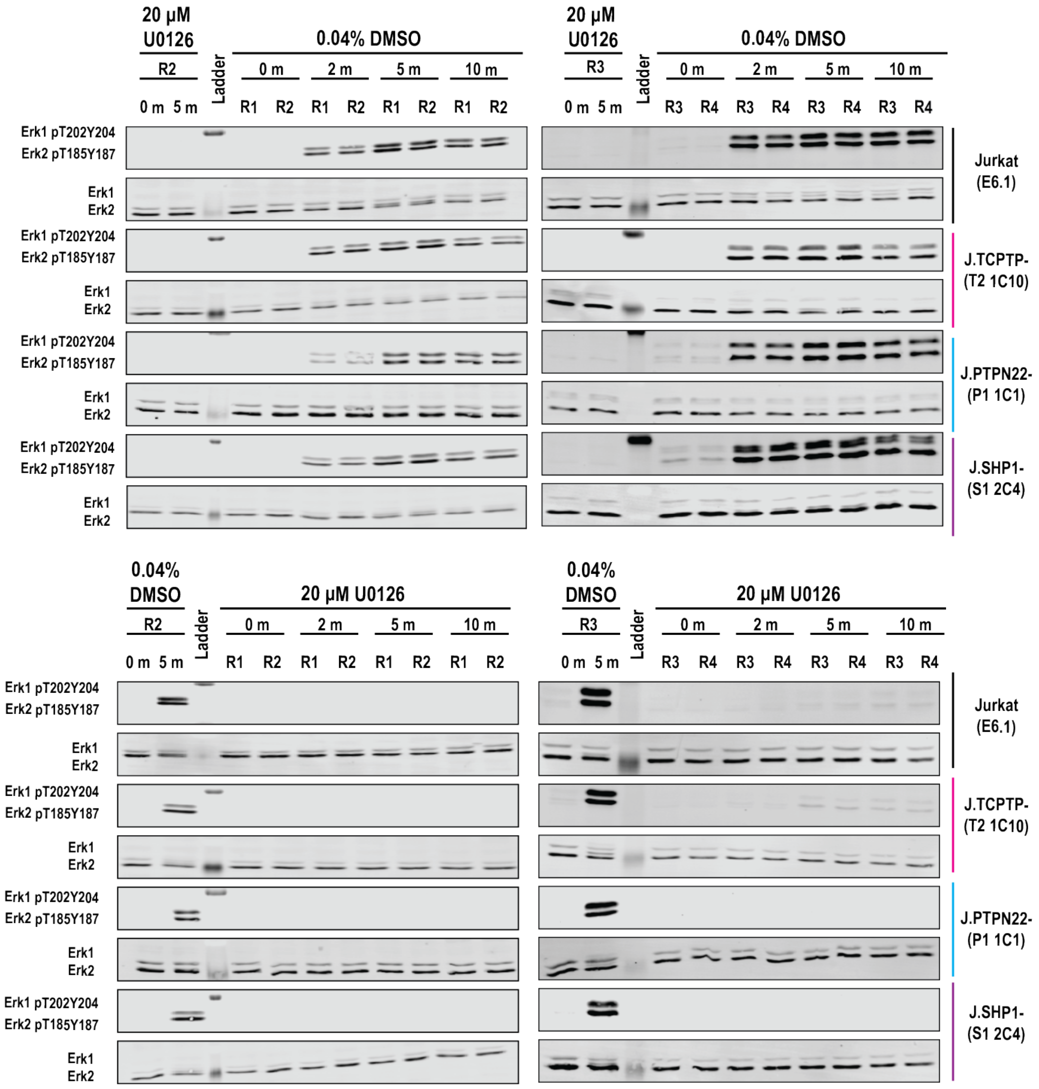


**Supporting Figure 4:** Western blots used for quantification of Erk1$^{T202Y204}$/Erk2$^{T185Y187}$ during a TCR stimulation time course in the absence (top) or presence (bottom) of 20 µM U0126.

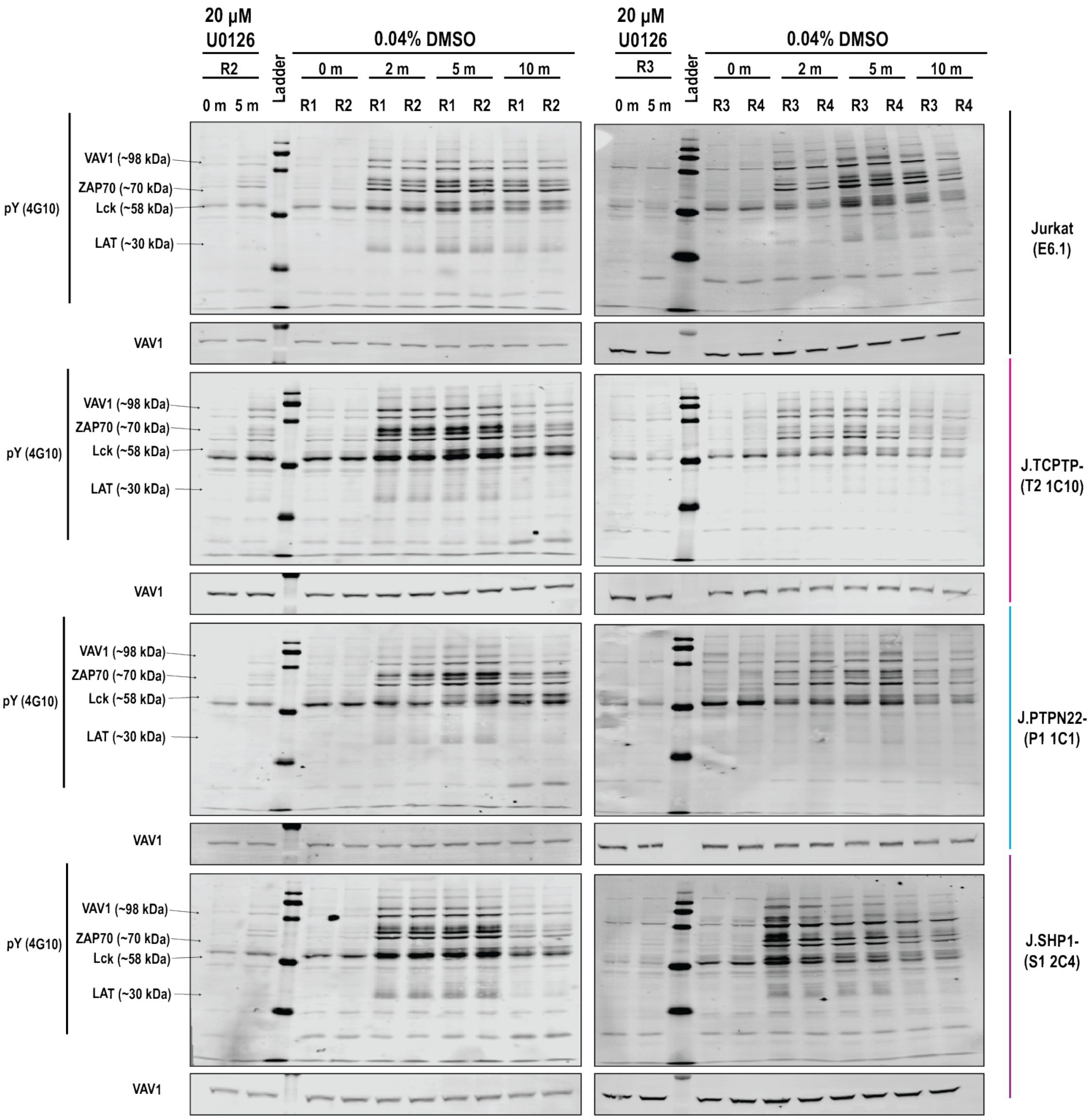


**Supporting Figure 5:** Western blots used for quantification of global pY abundance during a TCR stimulation time course in the presence of 0.04% DMSO.

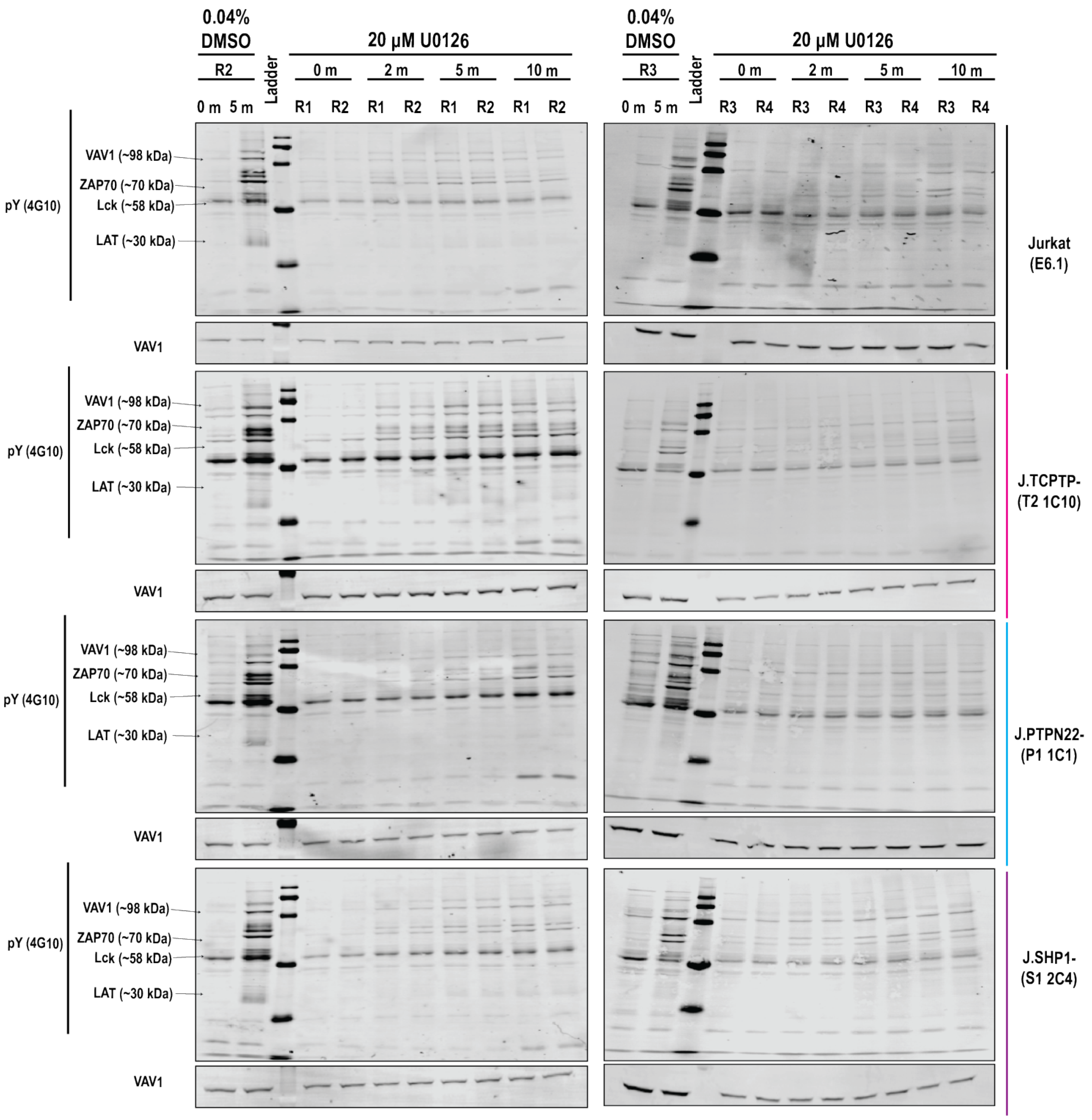


**Supporting Figure 6:** Western blots used for quantification of global pY abundance during a TCR stimulation time course in the presence of 20 µM U0126.

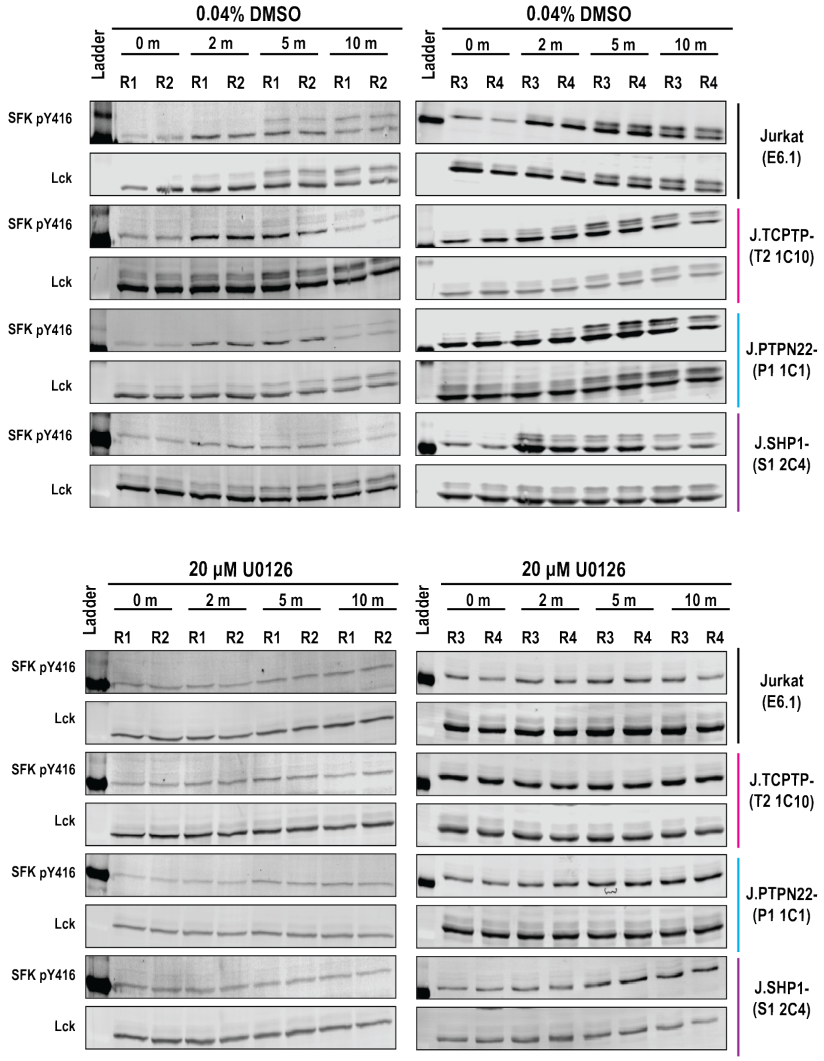


**Supporting Figure 7:** Western blots used for quantification of the Src family kinase activation site ($SFK^{Y416}$)during a TCR stimulation time course in the absence (top) or presence (bottom) of 20 μM U0126.

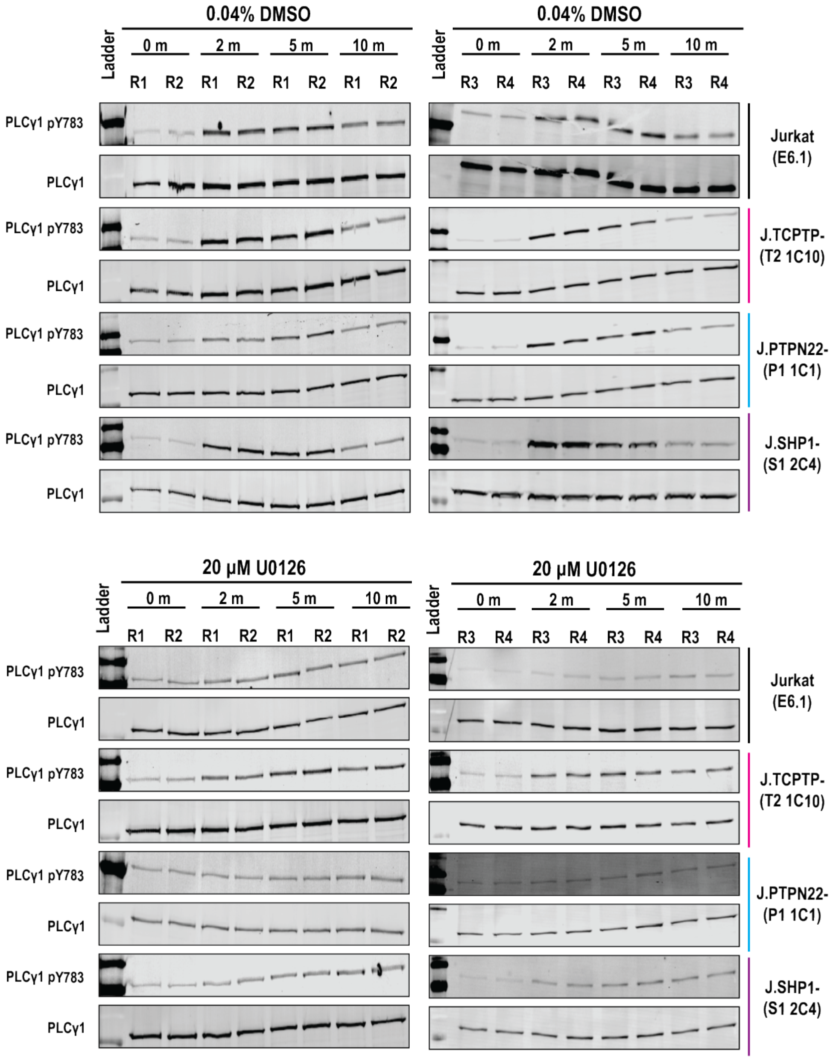


**Supporting Figure 8:** Western blots used for quantification of PLCγ1$^{Y783}$ during a TCR stimulation time course in the absence (top) or presence (bottom) of 20 µM U0126.

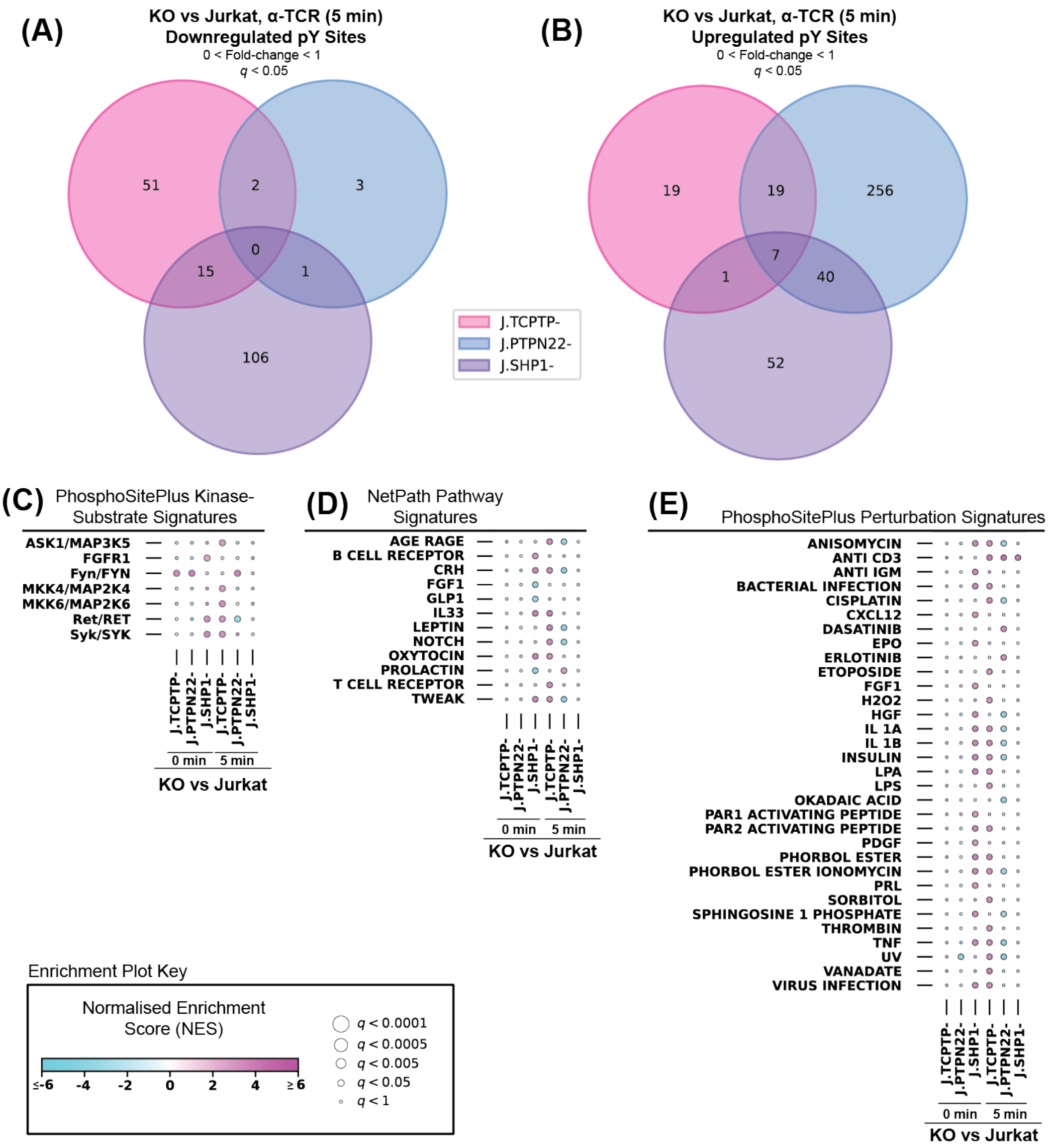


**Supporting Figure 9**: Comparison of significantly changing, unique pY sites and pathway enrichment in KOs compared to Jurkats. (A-B) Venn diagrams showing overlap of unique pY sites that are downregulated and upregulated, respectively, in J.TCPTP-, J.PTPN22-, and J.SHP1- compared with Jurkats during TCR stimulation. (C-E) Bubbleplots showing PTM-SEA of PhosphoSitePlus Kinase Signatures, NetPath Pathway Signatures, and PhosphoSitePlus Perturbation Signatures, respectively, for KO vs WT comparisons.

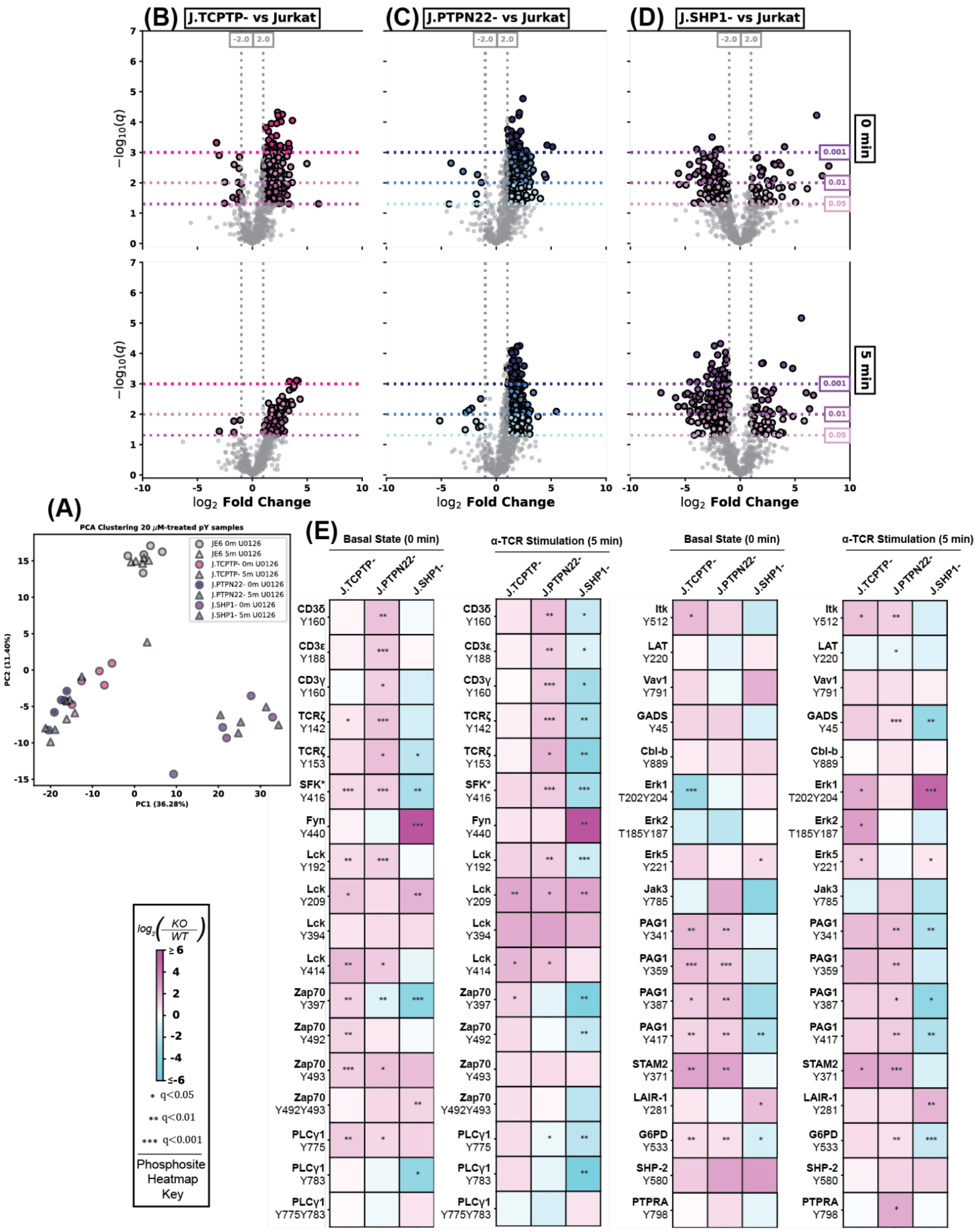


**Supporting Figure 10:** U0126 treatment does not change the distribution of pY abundance when comparing knockouts to Jurkats. (A) PCA showing clustering of samples treated with U0126. (B-D) Volcano plots comparing unique pY sites in J.TCPTP-, J.PTPN22-, and J.SHP1-, respectively, to Jurkats in the basal state (0 minute - top row) and stimulated state (5 minute - bottom row) during 20 µM U0126 treatment. (E) Heatmaps showing individual pY sites compared with Jurkats for each phosphatase KO during 20 µM U0126 treatment.

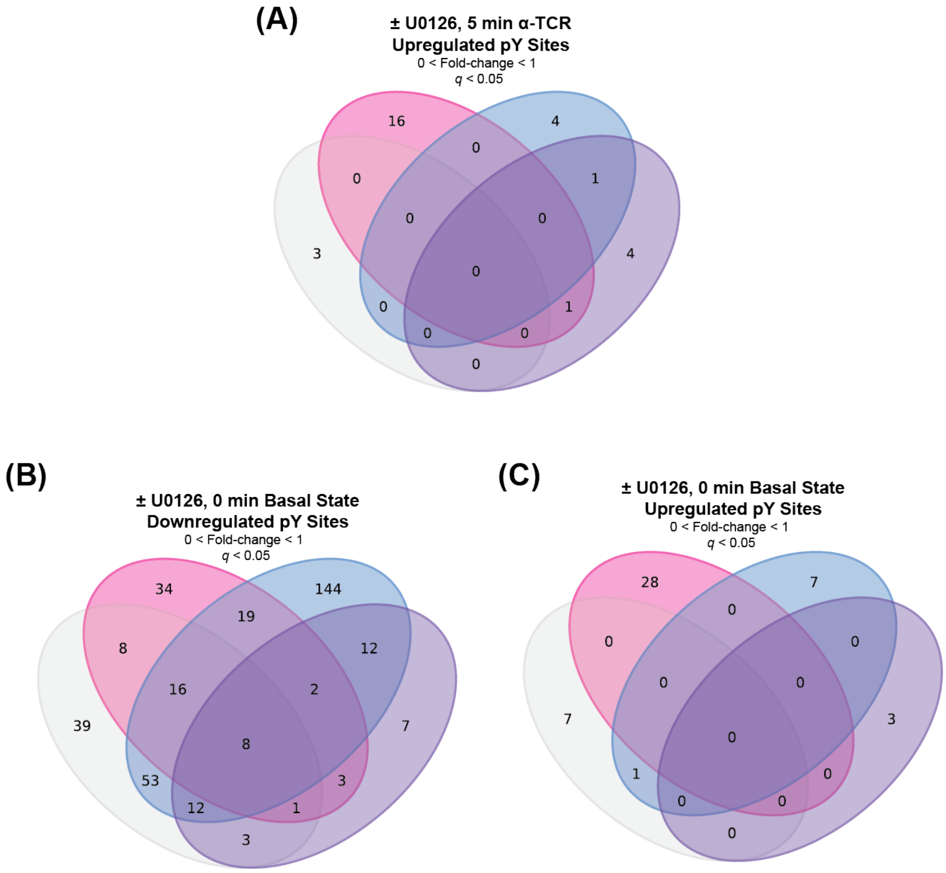


**Supporting Figure 11:** Effects of U0126 treatment on Jurkats, J.TCPTP-, J.PTPN22-, and J.SHP1-. (A) Venn diagram showing overlap of unique pY sites that are upregulated in response to U0126 treatment in the stimulated state. (B) Venn diagram showing overlap of unique pY sites that are downregulated in response to U0126 treatment in the basal state. (C) Venn diagram showing overlap of unique pY sites that are upregulated in response to U0126 treatment in the basal state.
